# The *Primula edelbergii S*-locus is an example of a jumping supergene

**DOI:** 10.1101/2024.01.30.577918

**Authors:** Giacomo Potente, Narjes Yousefi, Barbara Keller, Emiliano Mora-Carrera, Péter Szövényi, Elena Conti

## Abstract

Research on supergenes, non-recombining genomic regions housing tightly linked genes that control complex phenotypes, has gained prominence in genomics, with supergenes having been described in most eukaryotic lineages. Heterostyly, a floral heteromorphism promoting outcrossing in several angiosperm families, is controlled by the *S*-locus supergene. Historically, the *S*-locus has been studied primarily in closely related *Primula* species and, more recently, in other groups that independently evolved heterostyly. However, it remains unknown whether genetic architecture and composition of the *S*-locus are maintained among species that share a common origin of heterostyly and subsequently diverged across larger time scales. To address this research gap, we present a chromosome-scale genome assembly of *Primula edelbergii*, a species that shares the same origin of heterostyly with *Primula veris* (whose *S*-locus has been characterized) but diverged from it ca. 18 million years ago. Comparative genomic analyses between *P. edelbergii* and *P. veris* allowed us to show, for the first time, that the *S*-locus can ‘jump’ (i.e. translocate) between chromosomes. Additionally, we found that four *S*-locus genes were maintained across time but were reshuffled within the supergene, seemingly without affecting their expression. Furthermore, we confirmed that *S*-locus hemizygosity counteracts genetic degeneration, otherwise expected in supergenes. Finally, we investigated *P. edelbergii* evolutionary history within Ericales in terms of whole genome duplications and transposable element accumulation. In summary, our work provides a valuable resource for comparative analyses aimed at investigating the genetics of heterostyly and the pivotal role of supergenes in shaping the evolution of complex phenotypes.

## Introduction

Supergenes are non-recombining regions containing multiple, tightly linked loci, each controlling a different trait of the same complex phenotype (Thompson & Jiggins, 2014). Despite their prevalence across eukaryotic lineages (Schwander et al., 2014), supergenes have remained difficult to sequence until recently, owing to their distinctive molecular and evolutionary characteristics, as explained next. The lack of recombination among the genes in a supergene allows for the co-inheritance of only certain allelic combinations and the consequent occurrence of certain discrete phenotypes (Thompson & Jiggins, 2014). However, the lack of recombination is also expected to decrease the strength of purifying selection, with the consequent accumulation of deleterious mutations, transposable elements (TEs), deletions, and gene losses, in a process called genetic degeneration (Gutiérrez-Valencia et al., 2021). Because of these intrinsic features, supergenes are challenging regions to assemble, hence their study has long been precluded. However, recent advancements in genomic technologies have enabled the characterization of supergenes controlling wing-color patterning in butterflies (Joron et al., 2011), plumage and reproductive behavior in birds (Küpper et al., 2015; Lamichhaney et al., 2015; Tuttle et al., 2016), mating types in fungi (Branco et al., 2018), and reproductive features in plants (Li et al., 2016).

The first supergene ever described was the *S*-locus controlling heterostyly (Bateson & Gregory, 1905), a floral sexual heteromorphism consisting in the co-occurrence of two (distyly) or three (tristyly) types of flowers at the population level (Darwin, 1862). Heterostyly occurs in at least 28 angiosperm families and evolved independently at least twelve times (Ganders, 1979; Naiki, 2012) but, since Darwin, most of the research has focused on distylous primroses (i.e. species of the genus *Primula*, family Primulaceae, order Ericales; Darwin, 1862; Gilmartin, 2015). In *Primula*, two types of flowers coexist: pins (or long-styled (L)-morphs) have a long style and low anthers, while thrums (or short-styled (S)-morphs), have a short style and high anthers (**Figure 1a**). Furthermore, in most *Primula* species the floral dimorphism is associated with pollen and stigma papillae dimorphism (with thrums producing larger pollen grains and shorter papillae than pins), and an intra-morph incompatibility system that prevents fertilization between individuals of the same morph (Keller et al., 2014; Shivanna et al., 1981; Wedderburn & Richards, 1990). In *Primula*, the genetic architecture of heterostyly has been characterized only in two closely related species, *P. veris* and *P. vulgaris*, which diverged ca. 2.5 million years ago (Mya) (**Figure 1b**; de Vos et al., 2014a; Stubbs et al., 2022). In these two species the *S*-locus is ca. 260-kb long, contains the same five genes (*CCM*^*T*^, *GLO*^*T*^, *CYP*^*T*^, *PUM*^*T*^, and *KFB*^*T*^, hereafter collectively referred to as *S*-genes) in the same order, and occurs in the population as a presence-absence polymorphism, being hemizygous in thrums (*S*/*s*), and absent from pins (*s*/*s*) (Li et al., 2016; Potente et al., 2022a). Of the five *S*-genes, only two have been functionally characterized: *CYP*^*T*^ controls both style length and the female side of incompatibility (Huu et al., 2016, 2022), while *GLO*^*T*^ determines anther position (Huu et al., 2020).

**Figure 1:**
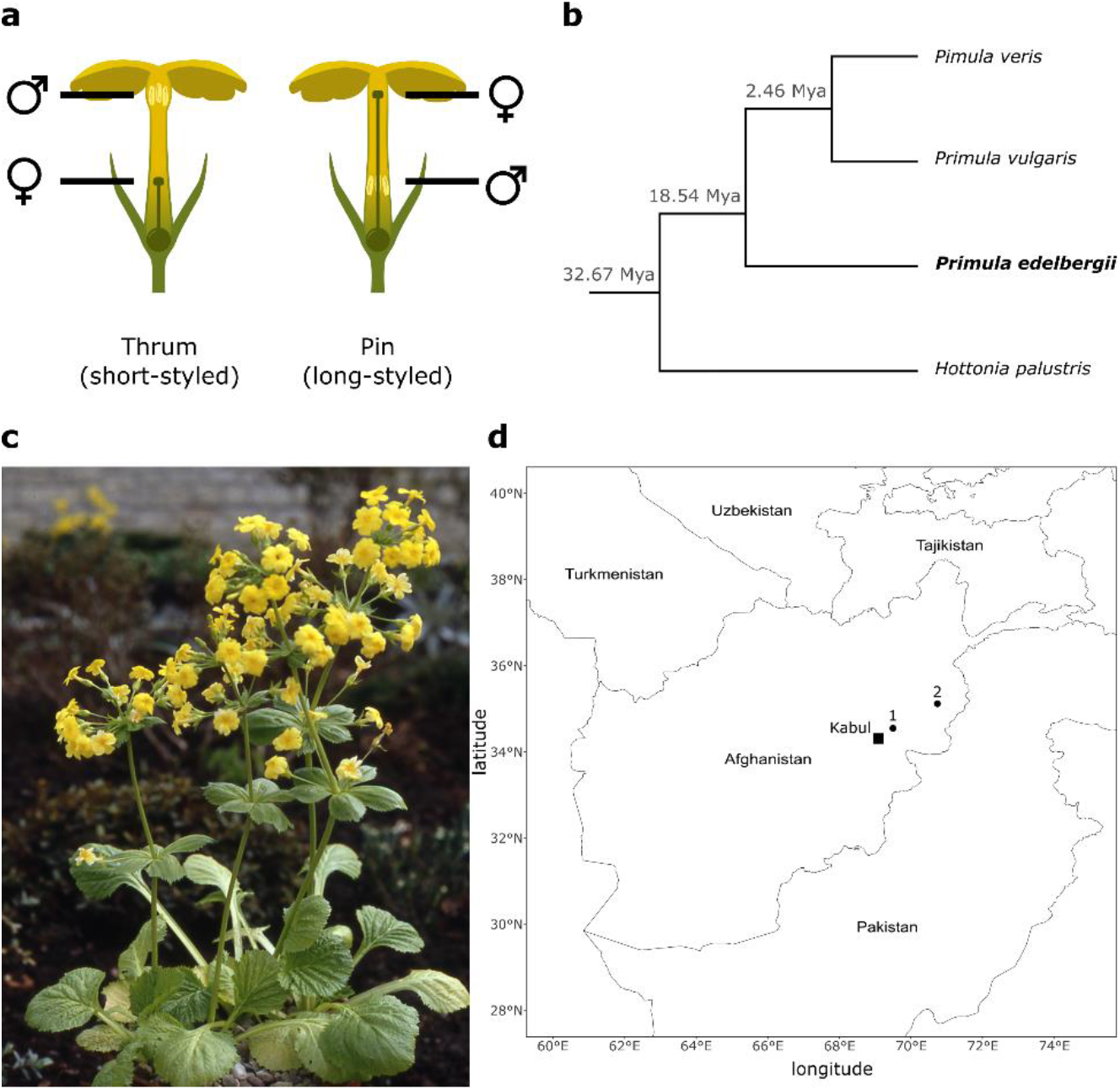
*Primula edelbergii* is an endemic primrose from Afghanistan. (**a**) Thrum (short-styled) and pin (long-styled) morphs of *Primula* species differ by having male (anthers) and female (stigma) sexual organs reciprocally positioned in their flowers. (**b**) Simplified phylogeny with node ages expressed in million years ago (Mya). The phylogeny includes: the two *Primula* species for which the *S*-locus has been sequenced and assembled (*P. veris* and *P. vulgaris*), which diverged ca. 2.46 Mya; *P. edelbergii*, which diverged from *P. veris* and *P. vulgaris* 18.54 Mya (de Vos, et al., 2014a); *Hottonia palustris*, sister to *Primula*, which diverged from it 32.67 Mya (The Angiosperm Phylogeny Group, 2016). (**c**) Picture of *P. edelbergii*. Credits to Prof. John Richards. (**d**) *P. edelbergii* is exclusively known from two sites in Afghanistan: the Tang-e Gharu gorge, in the Kabul province (1), and the Wama district, in the Nuristan Province (2).

Recently, research on the *S*-locus has expanded from *Primula* to other genera that independently evolved heterostyly, including *Turnera* (Henning et al., 2022; Matzke et al., 2020, 2021; Shore et al., 2019), *Fagopyrum* (Fawcett et al., 2023; Yasui et al., 2012, 2016), *Linum* (Gutiérrez-Valencia et al., 2022), *Gelsemium* (Zhao et al., 2023), and *Nymphoides* (Yang et al., 2023). Surprisingly, in all these cases the *S*-locus was found to be hemizygous in thrums and absent from pins, pointing to a remarkable consistency in the genomic architecture of the *S*-locus across heterostylous taxa. The recent characterization of the *S*-locus in distantly related families makes it an ideal model for investigating the convergent evolution of genetic architecture and its role in preserving complex polymorphisms. However, the genetic basis of heterostyly has been elucidated either in species stemming from independent origins of heterostyly, or closely related species such as *P. veris* and *P. vulgaris*, which diverged only ca. 2.5 Mya (**Figure 1b**; de Vos et al., 2014b; Stubbs et al., 2022) and exhibit such minimal genetic divergence that their respective *S*-loci are essentially identical (Cocker et al., 2018; Li et al., 2016; Potente et al., 2022a). Therefore, the extent of *S*-locus variation among species that diverged across larger timescales but share the same origin of heterostyly, encompassing aspects such as gene content, genomic location, genetic architecture, and degree of genetic degeneration, remains unknown.

To fill these gaps of knowledge, we generated a chromosome-scale genome assembly of *Primula edelbergii* (**Figure 1c**). We selected this species for four key reasons. First, *P. edelbergii* is distantly related to *P. veris* and *P. vulgaris*, as their most recent common ancestor was dated at ca. 18 Mya (de Vos et al., 2014a). Second, *P. edelbergii* is known from only two sites in eastern Afghanistan (**Figure 1d**; Richards, 2003) and, although an IUCN classification is lacking, we advocate that generating genomic resources for this species and investigating the genetics underlying its reproduction would be beneficial for its conservation both *in-situ* and *ex-situ*. Third, the karyotype of *P. edelbergii* consists of nine chromosomes (2n = 18; Richards, 2003), differing from *P. veris* and *P. vulgaris* (both 2n = 22). Therefore, assembling the genome of *P. edelbergii* would allow us to investigate which structural rearrangements led to the differential chromosome number among primroses. Fourth, *P. edelbergii* is self-compatible, while retaining the other key characteristics of heterostyly, i.e. reciprocal positioning of sexual organs and pollen dimorphism, thus representing a rare case among *Primula* species (Al Wadi & Richards, 1993; Wedderburn & Richards, 1990). Indeed, in all primroses studied so far (i.e. *P. veris, P. vulgaris, P. forbesii)*, both style length and the female side of incompatibility are controlled by the same *S*-gene, *CYP*^*T*^, and loss-of-function mutations in *CYP*^*T*^ lead to the formation of self-compatible flowers with stigma and anthers at the same height (Huu et al., 2016, 2022; Li et al., 2016; Mora-Carrera et al., 2021). Generating genomic resources for *P. edelbergii* could thus allow us to explain how the genetic control of style length can be decoupled from that of incompatibility.

Beyond heterostyly, *P. edelbergii* holds additional interest due to its classification within Ericales, an angiosperm order which contains several plants of economic interest, including crops such as blueberries (*Vaccinium*), kiwifruits (*Actinidia*), and tea (*Camellia*), and ornamentals such as primroses, cyclamens, and rhododendrons. Given their economic value, genome assemblies for multiple Ericales species have been published, enabling comparative genomics analyses. However, despite the keen interest in Ericales, their evolutionary history remains unclear, especially regarding the occurrence and timing of whole-genome duplications (WGDs; Larson et al., 2020; Nie et al., 2022). Multiple WGDs have been identified in Ericales, among which are *Ad-β*, a WDG at the root of Ericales (Larson et al., 2020; Shi et al., 2010), and *Pv-α*, a WGD detected in *P. veris* and *P. vulgaris* (Potente, et al., 2022). However, the absence of high-quality genome assemblies for any other Primulaceae species has so far hindered our ability to determine whether Pv-α was specific to *Primula* and whether other WGDs occurred in Primulaceae.

Here, we present a chromosome-scale genome assembly for *P. edelbergii*. Such a high-quality genomic resource, analyzed in comparison with the genomes of other Ericales, enabled the following key discoveries. First, the *S*-locus is not in the same genomic position in *P. edelbergii* and *P. veris*, indicating that the entire supergene was translocated in one of the species since their divergence. Secondly, the *S*-locus of *P. edelbergii* comprises only four of the five *S*-genes found in *P. veris* and *P. vulgaris*, and they occur in a different order in *P. edelbergii*. Third, we detected no signatures of genetic degeneration in the S-locus of *P. edelbergii* and *P. veris*. Our study provides, to our knowledge, the first evidence for the translocation of a supergene not related to sex determination and represents a useful genomic resource for further comparative genomic analyses of heterostyly and, more broadly, supergenes. Finally, the *P. edelbergii* genome represents a new resource to discover the genomic processes driving chromosome-number evolution in angiosperms.

## Materials and Methods

### Plant material

The genome plant was a thrum individual (**Figure 1a**) grown from seeds that we obtained from the Botanical Garden München-Nymphenburg (Germany; XX-0-Z-20050490), an accession that was originally collected by Per Wendelbo in Afghanistan in 1969 (collection no. PW 9739). To our knowledge, all plants currently cultivated at botanical gardens are thrums and originate from the Wendelbo collection.

### Nucleic acid extraction and sequencing

High molecular weight DNA was extracted from fresh leaves following a protocol we previously developed (Potente et al., 2022a). After DNA extraction, secondary plant compounds hampering the activity of the pores in the Oxford Nanopore Technologies (ONT) flow cells were removed with the Quick-DNA HMW MagBead Kit (Zymo Research). Sequencing libraries were prepared with the LSK-109 ligation library preparation kit and sequencing was performed on MinION R.9.4.1 flow cells. Basecalling was done using Guppy v6.0.1 and the Super High Accuracy (SUP) model, resulting in 2.84 million reads (39.79 Gb, corresponding to 59x coverage; read N50 = 19.16 kb). We also generated 545.60 million paired-end (PE) 151-bp reads (Illumina NovaSeq; Functional Genomic Center Zurich (FGCZ), Switzerland), totaling 82.39 Gb (123x coverage). Hi-C libraries for scaffolding were generated and sequenced by Arima Genomics (200.48 million PE 150-bp reads; Illumina NovaSeq; Arima Genomics, USA). To aid gene annotation, we extracted RNA from leaves, young floral buds (i.e. petals did not protrude the calyx), mature floral buds (i.e. petals protruded the calyx, but were still closed), and full blooming flowers (i.e. petals were fully unfolded) using the Spectrum Plant Total RNA Kit (Sigma Aldrich) and generated 80.00 million PE 150-bp reads (Illumina NovaSeq; FGCZ, Switzerland).

### Genome-size estimate, genome profiling, and genome assembly

We estimated the genome size of *P. edelbergii* using *k*-mers. Specifically, we used half of the Illumina dataset (i.e. the forward reads) to count 21-mers with the *count* function of Jellyfish v2.2.10 (-C -m 21; Marçais & Kingsford, 2011). Then we used the *histo* function of Jellyfish v2.2.10 to generate a suitable input file for the online version of GenomeScope (*k*-mer length: 21; read length: 150; max *k*-mer coverage: 10,000; <u>qb.cshl.edu/genomescope</u>; Vurture et al., 2017).

Illumina and ONT reads were used to assemble the *P. edelbergii* genome using the hybrid approach in MaSuRCA v4.0.5 (Zimin et al., 2017), with Flye v2.5 (Kolmogorov et al., 2019) for the final “contigging” step. The resulting assembly (671.80 Mb; N50 = 12.42 Mb; L50 = 17) was scaffolded using Hi-C reads in YaHS v1.2a.1 (C. Zhou et al., 2023), producing an assembly of 671.76 Mb, with N50 = 73.57 Mb and L50 = 3. We generated a Hi-C contact map and visualized it in Juicebox (JBAT; Dudchenko et al., 2018) for manual curation. The final *P. edelbergii* genome assembly was 671.82 Mb long (N50=71.63 Mb; L50=5). The quality of the genome assembly was evaluated using the Kmer Analysis Toolkit (KAT) v2.4.2 (Mapleson et al., 2017), which required a *k*-mer ‘histo’ file generated with Jellyfish v2.2.10 as described above. Finally, genome completeness was evaluated with BUSCO v4.0.6 (Manni et al., 2021).

### Gene annotation

Gene annotation was performed on the *P. edelbergii* genome using a combination of *ab initio* and homology-based methods by providing BRAKER v3.0.1 (Brůna et al., 2021) with RNA-seq data from four tissues (leaf, flower, young floral bud, mature floral bud) and protein sequences from species of “unknown evolutionary distance”. First, RNA-seq reads were trimmed using Trimmomatic v0.38 (Bolger et al., 2014), with the parameters recommended by Trinity (Grabherr et al., 2011): ILLUMINACLIP:2:30:10 SLIDINGWINDOW:4:5 LEADING:5 TRAILING:5 MINLEN:25. Trimmed RNA-seq reads were then aligned against the *P. edelbergii* genome assembly (soft-masked as described below using RepeatMasker v4.0.9; www.repeatmasker.org) using Hisat v2.1.0 (--dta --max-intronlen 50000 --fr; Kim et al., 2019). Second, a set of protein sequences for homology-based annotation was prepared by merging the OrthoDB protein data set for Viridiplantae (odb10; www.orthodb.org) with a high-quality *P. veris* protein set. Such high-quality *P. veris* protein set was newly created here by filtering, from the entire *P. veris* protein set, only those proteins that a) were encoded by genes that had two, or more, exons (because the *P. veris* annotation was done using an older version of BRAKER, known to generate an excess of monoexonic gene models; Vuruputoor et al., 2023) and b) were supported by RNA-seq evidence for their entire length (Potente et al., 2022a). The final protein set consisted of 3,529,642 proteins (18,900 of which were *P. veris* proteins). Finally, BRAKER3 v3.0.1 (Brůna et al., 2021) was run on the *P. edelbergii* genome assembly with default parameters, providing as input the protein set consisting of the OrthoDB set merged with the high-quality *P. veris* protein set and the RNA-seq alignment file.

We assessed the quality of gene annotation in two ways. First, the percentage of each gene model covered by RNA-seq was assessed by running the ERE and AnnotationEvidence functions of GeMoMa v1.6.2 (Keilwagen et al., 2016, 2018). Second, BUSCO v4.0.6 (Manni et al., 2021) was run on the coding sequences of *P. edelbergii* and nine other Ericales species (see below), using the 2,326 single-copy orthologs from the eudicot database (odb v10).

### Quantification of gene expression

We quantified gene expression from four tissues (see above). To determine the possible causes for the lack of self-incompatibility in *P. edelbergii*, we investigated gene expression in S-genes. Trimmed RNA-seq reads from the four tissues were used to quantify expression of each transcript with the Salmon v1.4.0 (Patro et al., 2017) *quant* function (mapping-based mode; –gcBias –validateMappings). Salmon output was then imported into R v4.1.2 (www.R-project.org/) using tximport (Love et al., 2018; Soneson et al., 2016) and a DESeqDataSet was created with the DESeqDataSetFromTximport function of the DESeq2 (Love et al., 2014) R/Bioconductor (Gentleman et al., 2004) package. At this stage, expression was summarized at the gene level. Gene counts were then normalized using the default median of ratios method (Love et al., 2014). Finally, we classified each gene as expressed or not in each tissue. To do so, we first plotted the distribution of normalized RNA-seq counts for all genes in each tissue, which resulted in a distribution in which only the main peak at higher expression (centered at ca. 100 normalized reads; **Figure 3a**) includes the active, functional transcriptome, i.e. expressed genes (Hebenstreit et al., 2011). Thus, following Hart et al., 2013, we marked a gene as expressed in a tissue if its normalized RNA-seq count belonged to the peak at higher expression by setting a conservative threshold of normalized RNA-seq reads >= 16, as done for *P. veris* in a previous study (Potente et al., 2022b).

**Figure 2:**
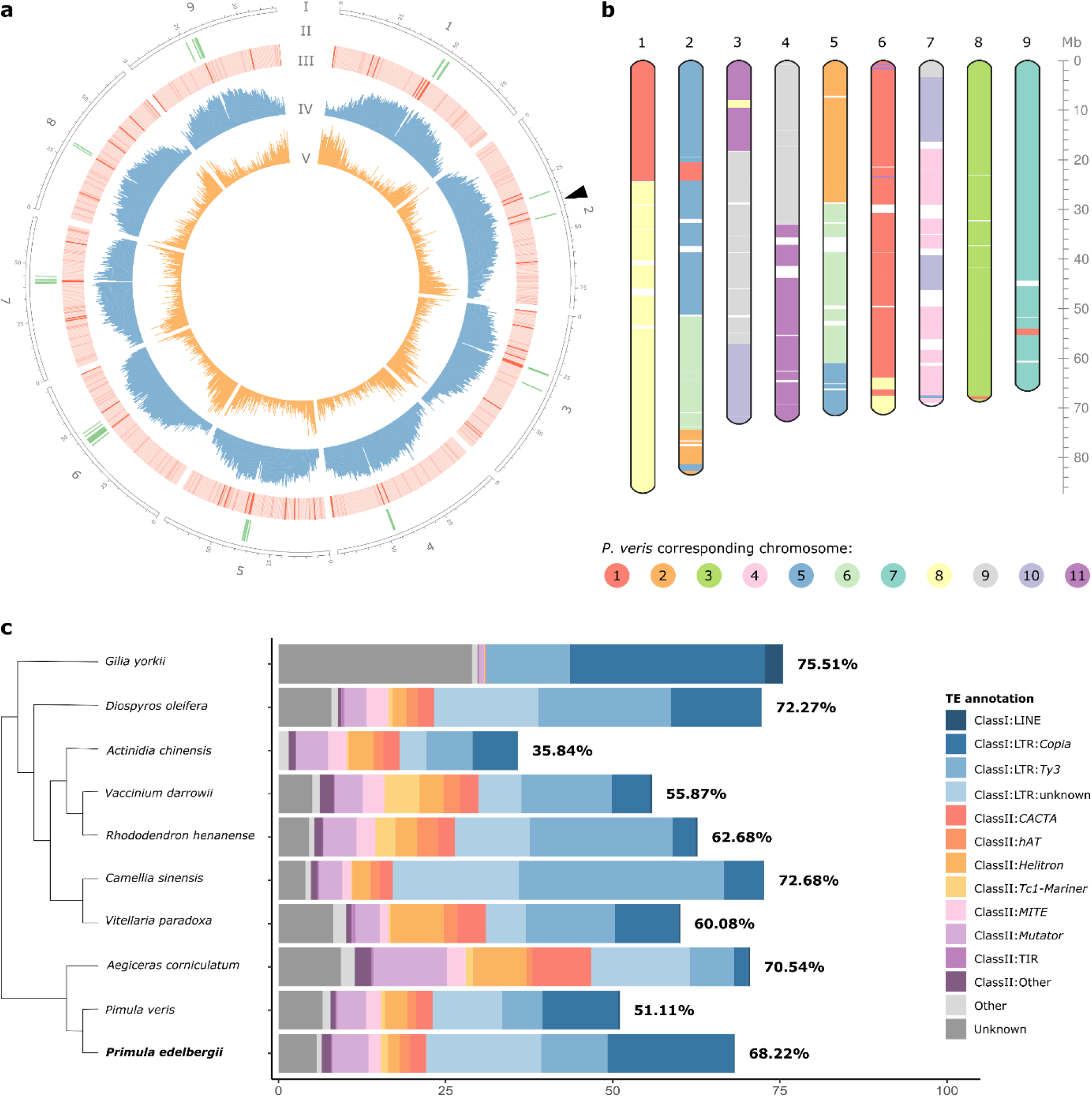
Summary of the *P. edelbergii* genome assembly and comparison with *P. veris*. (**a**) Circle plot of the *P. edelbergii* genome assembly. Tracks from outside to inside correspond to: (I) the nine chromosomes, with the position of the *S*-locus marked by a black arrow in chromosome 2; (II) BLAST matches of the putative centromeric tandem repeats (green); (III) GC density, with darker red bars indicating higher GC content; (IV) DNA transposons density (blue); (V) gene density (orange). Tracks II to V were calculated in 500-kb non-overlapping windows. (**b**) Chromosome-painting plot representing the nine *P. edelbergii* chromosomes, with regions syntenic to different *P. veris* chromosomes colored as indicated by the legend. Genomic regions that are not syntenic between the two species are shown in white. (**c**) Whole-genome TE abundance, expressed as percentage of genome covered by each TE superfamily as in legend, for the ten Ericales species analyzed here. The cladogram on the left shows phylogenetic relationship of the species.

**Figure 3:**
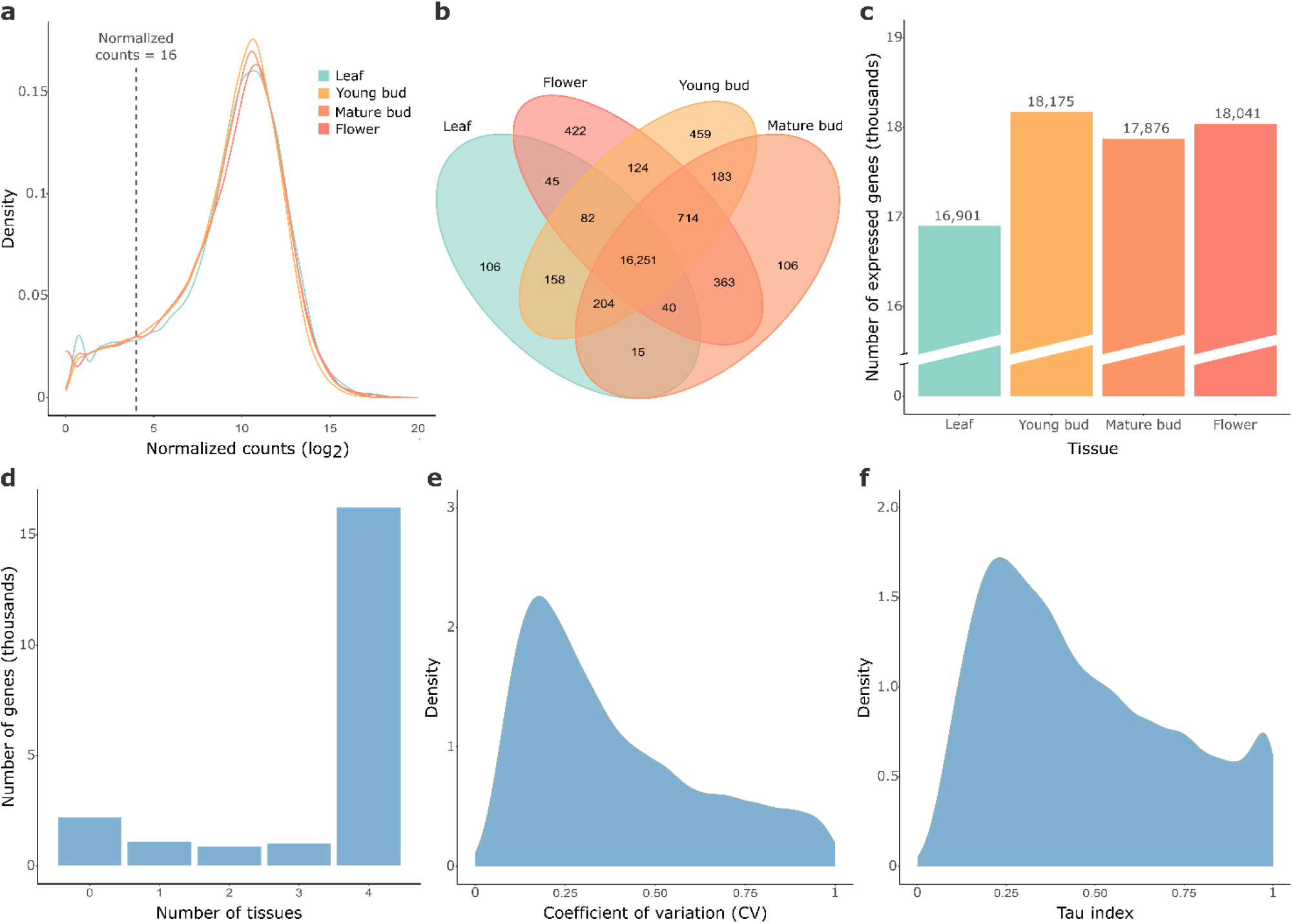
Overview of the *P. edelbergii* transcriptome. (**a**) Density plot showing the distribution of normalized RNA-seq read counts for all genes, with each line representing a tissue, as indicated in the legend. The vertical dashed line indicates the threshold above which we marked a gene as expressed in a tissue. (**b**) Venn diagram indicating the number of expressed genes shared by each tissue combination; 16,251 genes (75.7% of the total) are expressed in all tissues. (**c**) Number of expressed genes in each tissue; floral tissues (warm colors) are characterized by a higher number of expressed genes than the leaf tissue (turquois). (**d**) Number of genes expressed in any given number of tissues; most genes are expressed in all four tissues. (**e, f**) Density plots showing the distribution of two tissue-specificity metrics across all genes, i.e. the coefficient of variation (CV; e), and the Tau index (f).

### Comparative genomics analyses across Ericales

We compared the genome of *P. edelbergii* with those of other species of the same order (Ericales) to investigate the evolution of TEs (**Figure 2c**). We selected species for which a chromosome-scale genome assembly was available, with the aim of covering as many Ericales families as possible. For our comparative genomics analyses we used the genome assemblies of ten Ericales species, representing seven of the 22 Ericales families (The Angiosperm Phylogeny Group, 2016), including: *P. edelbergii* (present study), *P. veris* (Potente et al., 2022a), and *Aegiceras corniculatum* (Ma et al., 2021; Primulaceae); *Actinidia chinensis* (Pilkington et al., 2018; Actinidiaceae); *Rhododendron henanense* (X. J. Zhou et al., 2022) and *Vaccinium darrowii* (Yu et al., 2021; Ericaceae, subfamilies Ericoideae and Vaccinioideae, respectively); *Diospyros oleifera* (Suo et al., 2020; Ebenaceae); *Vitellaria paradoxa* (Hale et al., 2021; Sapotaceae); *Camellia sinensis* (Wei et al., 2018; Theaceae); *Gilia yorkii* (Jarvis et al., 2022; Polemoniaceae).

### Repeat annotation

Repetitive elements were identified in the genome assemblies of all Ericales species analyzed for this study (excluding *Gilia yorkii*, see below), as follows. Extensive De novo TE Annotator (EDTA) v1.9.2 (Ou et al., 2019) was run on each genome assembly separately to identify repetitive sequences and build a consensus repeat library for each species. EDTA is a pipeline that combines structure- and homology-based approaches for *de novo* TE identification. Structure-based TE discovery was performed using LTRharvest (Ellinghaus et al., 2008) and LTR_retriever (Ou & Jiang, 2018) for long terminal repeats retrotransposons (LTR-RTs), TIR-Learner (Su et al., 2019) for terminal inverted repeats (TIR) elements, and Helitronscanner (Xiong et al., 2014) for helitrons. Other repetitive elements were identified using RepeatModeler v2.0.1 (Flynn et al., 2020). For *Gilia yorkii*, repetitive elements were identified using RepeatModeler v2.0.1 (Flynn et al., 2020). Finally, each repeat library was used to annotate the respective genome assembly using RepeatMasker v4.0.9 (www.repeatmasker.org).

### Investigation of potential hemizygosity and heterozygosity in the *S*-locus

We identified the *S*-locus in *P. edelbergii* by mapping the protein sequences of the *P. veris S*-genes against the proteome of *P. edelbergii* using BLASTp (-evalue 1e-5; Camacho et al., 2009) The *P. edelbergii S*-genes are: g4960 (*CYP*^*T*^), g4963 (*PUM*^*T*^), g4964 (*GLO*^*T*^), g4966 (*KFB*^*T*^); no homolog was found for *CCM*^*T*^. To investigate the potential hemizygosity of the *S*-locus, we searched whether this region was characterized by a halved sequencing depth compared to the rest of the genome. We estimated sequencing depth across the *S*-locus and its flanking regions based on a VCF obtained by aligning short reads of the same individual used to generate the assembly to the *P. edelbergii* reference genome. Illumina reads were trimmed using Trimmomatic v0.38 (Bolger et al., 2014) with the following parameters: LEADING:3 TRAILING:3 SLIDINGWINDOW:4:15 MINLEN:36. Trimmed reads were then mapped to the *P. edelbergii* genome assembly using BWA-MEM v0.7.17 (H. Li, 2013) and the alignment file was sorted, marked for duplicates, and indexed using Picard (<u>broadinstitute.github.io/picard/</u>) and SAMtools v.1.9-63 (H. Li, 2011). The indexed alignment file was used as input for variant calling with the mpileup function of BCFtools v1.8 (Danecek et al., 2021; H. Li, 2011). After masking repetitive elements (identified as described above), we performed a filtering step with VCFtools v.0.1.14 (Danecek et al., 2011; --max-meanDP 210; --minQ 200). To generate the coverage plot of **Figure 4d** (top panel) we first extracted information on sequencing depth from the final VCF and then calculated depth in sliding windows of 10,000 sites (step = 1 site) using the rollmean function of the zoo package v1.8.12 (Zeileis & Grothendieck, 2005) in R v4.1.2 (www.R-project.org/). The final VCF was also filtered to keep only heterozygous SNPs, and their position was plotted in **Figure 4d** (bottom panel).

**Figure 4:**
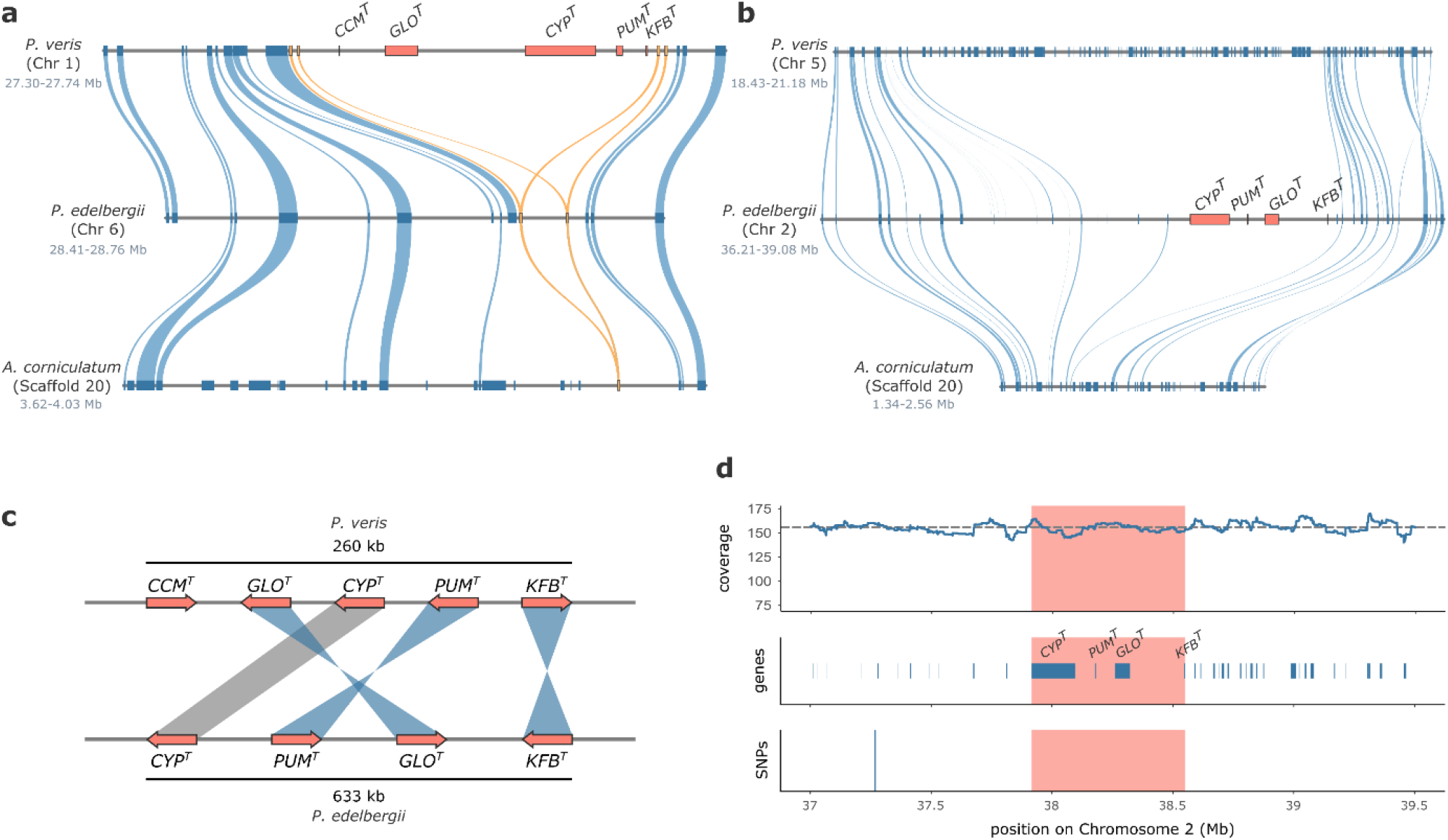
The *S*-locus translocated and is homozygous in *P. edelbergii*. (**a**) Microsynteny plot showing the region of chromosome 1 of *P. veris* containing the *S*-locus flanked by two *CFB* copies on each side (top), its syntenic region in chromosome 6 of *P. edelbergii* containing no *S*-gene and only two *CFB* copies (middle), and its syntenic region in *A. corniculatum* containing only one *CFB* copy (bottom). *S*-genes are represented as red boxes, *CFB* genes as yellow boxes, and other genes as blue boxes. Blue ribbons connect homologous gene pairs and yellow ribbons connect *CFB* homologs. (**b**) Microsynteny plot showing the region of chromosome 2 of *P. edelbergii* containing the *S*-locus (middle) and its syntenic regions in chromosome 5 of *P. veris* (top) and in scaffold 20 of *A. corniculatum* (bottom). Genes and ribbons are colored as in (a). (**c**) Schematic view of the *S*-locus structure in *P. veris* (top) and *P. edelbergii* (bottom), with *S*-genes represented as red arrows, whose directions indicate gene orientations. Blue ribbons connect orthologous *S*-genes whose orientations differ between the two species; a grey ribbon connects the two orthologous copies of *CYP*^*T*^, whose orientation remained the same. (**d**) Sequencing coverage (top), gene (middle) and SNP (bottom) positions in the *S*-locus (red boxes) and its flanking regions in *P. edelbergii* chromosome 2 (37.0-39.5 Mb). Top: the sequencing coverage (calculated in 10,000-SNPs sliding windows) is shown as a blue line; a grey dashed line indicates the mean coverage across chromosome 2. Middle: genes are shown as blue boxes, with names written on *S*-genes. Bottom: the only SNP between the two haplotypes in this region is indicated by a vertical dark blue line at position 37,265,444, located in an intergenic region.

### Investigation of *S*-locus degeneration in *P. edelbergii* and *P. veris*

To test for TE accumulation in the *S*-locus, we calculated the fraction of sequence occupied by TEs in 25-kb non-overlapping windows across the whole genomes of *P. edelbergii* and *P. veris*. We then subset the windows in three categories. First, the windows contained in the *S*-locus (n=27 for *P. edelbergii*; n=11 for *P. veris*). Second, the regions flanking the *S*-locus, which for *P. edelbergii* were regions containing the same number of 25-kb windows as the *S*-locus in total (n=27, including 13 windows downstream and 14 windows upstream), and for *P. veris* were the same number of 25-kb windows as the *S*-locus on each side (n=22, including 11 windows downstream and 11 windows upstream for *P. veris*). Third, the remaining windows were considered as the genomic background (n=26,556 for *P. edelbergii*; n=15,858 for *P. veris*). A Wilcoxon rank-sum test was used to test for significant differences between the three distributions in each species. To compare gene density within the *S*-locus to the rest of the genomes, we generated for each species a null gene-density distribution by randomly sampling 10,000 windows as long as the *S*-locus, i.e. 633,644 bp for *P. edelbergii* and 266,471 bp for *P. veris*, and counted the number of genes in each window. To estimate the strength of purifying selection on *S*-locus genes in *P. edelbergii* and *P. veris* we calculated non-synonymous substitutions per non-synonymous site (d_N_) and synonymous substitutions per synonymous site (d_S_) for all genes in the two genomes using *P. vulgaris* (Cocker et al., 2018) for comparison. First, we identified 12,827 orthologous gene pairs between *P. edelbergii* and *P. vulgaris* and 11,132 between *P. veris* and *P. vulgaris* using Proteinortho v6.0.31 (-p=blastn; Lechner et al., 2011). Then, d_N_ and d_S_ were calculated using ParaAT v2.0 (Zhang et al., 2012), which employs MUSCLE v3.8.31 (Edgar, 2004) to align sequences and KaKs_Calculator v2.0 (Wang et al., 2010) to calculate d_N_ and d_S_.

### Synteny analyses and WGD identification

Syntenic regions were identified within and among *P. veris, P. edelbergii* and *A. corniculatum* using MCScan (github.com/tanghaibao/jcvi/wiki/MCscan-(Python-version); Tang et al., 2008). We then estimated d_S_ using ParaAT v2.0 as explained above. For each d_S_ distribution, we computed median and standard deviation of statistically significant peaks by fitting a mixture of 1–3 normal distributions to the d_S_ histogram using R scripts (github.com/gtiley/Ks_plots; Tiley et al., 2018).

The d_S_ values calculated within *P. edelbergii* and *A. corniculatum* were used to infer WGDs, while the d_S_ value calculated between *P. edelbergii* and *A. corniculatum* was used to estimate the divergence time between the two species. To infer absolute ages for WGDs and species divergence, we used the neutral substitution rate of 6.15 × 10^−9^ substitutions per synonymous site per year previously determined for *P. veris* (Potente et al., 2022a). To account for potential differences in the accumulation of synonymous substitutions among species, we estimated a neutral substitution rate also for *A. corniculatum*. To do so, we first estimated d_S_ between *A. corniculatum* and *Actinidia chinensis* as d_S_ = 1.16; then, based on a previously reported divergence time between the two species of 97.6 Mya (Foster et al., 2017; Rose et al., 2018), we estimated a neutral substitution rate of 5.94 × 10^−9^ substitutions/year with the formula *r=d*_*S*_*/2t* (where *r* is the neutral substitution rate and *t* is the divergence time expressed in years). Using this substitution rate, the main conclusions did not change, as the putative WGD in *A. corniculatum* still resulted too recent to correspond to *Pv-α* (51.77 Mya; **Table S8**).

## Results and Discussion

### Overview of the *P. edelbergii* genome assembly and comparison with *P. veris*

Using a combination of long (nanopore) and short (Illumina) reads, plus Hi-C scaffolding, we generated a *P. edelbergii* genome assembly of 671.82 Mb (N50 = 71.63 Mb). The final assembly size aligns with the estimate obtained via *k*-mer analysis (**Figure S1**) and the nine largest scaffolds, ranging from 66.66 Mb to 87.23 Mb and totalizing 99.00% of the assembly, correspond to the nine chromosomes of the *P. edelbergii* haploid karyotype (**Figure 2a; Table 1**; **Table S1**). The high quality and completeness of the assembly was confirmed by *k*-mer (**Figure S2**) and BUSCO analyses (92.9% complete genes; **Figure S3, Table S2**).

**Table 1:** statistics for the *P. edelbergii* genome assembly and annotation.

|  |  |
| --- | --- |
| Assembly size | 671.83 Mb |
| Number of scaffolds | 376 |
| Cumulative length of the nine largest scaffolds | 665.10 Mb |
| Proportion of the assembly in the nine largest scaffolds | 99.00% |
| N50 | 71.63 Mb |
| L50 | 5 |
| BUSCO complete genes (genome) | 92.90% |
| no. genes | 21,513 |
| BUSCO complete genes (proteome) | 95.50% |
| genome fraction covered by TEs | 68.22% |

The central region of each chromosome showed a narrow window depleted in transposable elements (TEs) and enriched in GC content (**Figure 2a**). We investigated whether such regions corresponded to centromeric repeats. We extracted the sequence from one of these low-TE, high-GC regions and aligned it against itself, revealing that it was composed of a 773-bp motif repeated in tandem. By aligning this motif onto the *P. edelbergii* assembly, we determined that all nine low-TE, high-GC regions were composed by the same 773-bp motif, indicating that they are indeed centromeric repeats (**Fig. 2a**; **Figure S4**).

As the genome of *P. veris* comprises eleven chromosomes and that of *P. edelbergii* only nine, we investigated the chromosomal rearrangements that could explain this interspecific difference in chromosome number. Specifically, we asked whether the difference could be attributed solely to two chromosomal fusion/fission events or more extensive rearrangements had occurred. Synteny analyses revealed that multiple chromosomal rearrangements occurred since the divergence of these two species (**Figure 2b**; **Figure S5**). Notably, each *P. edelbergii* chromosome displayed synteny with at least two *P. veris* chromosomes, except for chromosomes 8 and 9 which, despite a few inversions, exhibited synteny with *P. veris* chromosomes 3 and 7, respectively. These findings suggest that several chromosomal translocations occurred since the divergence between the two species ca. 18 Mya (de Vos et al., 2014a). Despite the importance of chromosomal rearrangements in adaptation and speciation, little is known on how frequently they occur in different species (Lucek et al., 2023; Mérot et al., 2020), thus it is impossible to compare the rate of rearrangements observed in *Primula* with other plant groups. However, we previously discovered a high frequency of intra-species structural rearrangements in *P. veris* and postulated that this phenomenon may be linked to the dynamic nature of *Primula* genomes, potentially influenced by transposable elements (TEs), which constitute a significant portion of primrose genomes (see below; Potente et al., 2022a).

### The difference in genome size between *P. edelbergii* and *P. veris* is due to TE accumulation

To characterize the repetitive content of the *P. edelbergii* genome and compare it with related species, we estimated the abundance of different TE superfamilies in ten Ericales genomes. Transposable elements comprised on average 55.63% of the genome size in all Ericales species (min = 35.84%; max = 75.51%) and 68.22% in *P. edelbergii* (**Figure 2c**; **Table S3**; **Table S4**).

Given that *P. edelbergii* has a genome 1.6 times larger than *P. veris* (671.83 Mb vs 421.38 Mb), we investigated whether such a difference could be explained by differential TE accumulation. Indeed, we found that the overall length of non-repetitive genomic content of *P. edelbergii* was roughly the same as that of *P. veris* (213.51 Mb and 206.02 Mb, respectively) and the difference in genome size between the two species was explained by a 2.13 times higher content of repetitive elements in *P. edelbergii* (458.32 Mb, corresponding to 68.22% of the genome size) than in *P. veris* (215.36 Mb, 51.11% of the genome size). Specifically, *P. edelbergii* showed an increase in 19 of the 23 TE superfamilies considered, with each TE superfamily being on average 1.86 times more abundant in *P. edelbergii* than *P. veris*. However, differences were found in the relative abundance of different TE classes. *Primula edelbergii* and *P. veris* had a similar amount of Class I (DNA) transposons, which covered 13.54% and 12.91% of the two genomes, respectively, but different superfamilies showed different patterns: *hAT, CACTA*, and *Mutator* were 1.77 times more abundant in *P. edelbergii* than in *P. veris*, concordantly with the average for all TE superfamilies; *PIF-Harbinger* and *Tc1-Mariner* were over-represented in *P. edelbergii*, being 2.83 and 2.67 times more abundant than in *P. veris*; finally, *Helitron* were more abundant in *P. veris* than in *P. edelbergii*. Unlike DNA transposons, (Class II) LTR-RT content differed greatly between the two species, accounting for 46.01% of *P. edelbergii* genome size (309.19 Mb) but only 27.73% of *P. veris* genome size (116.83 Mb; **Figure 2c**), therefore representing the main cause of differential genome size between the two species. Among LTR-RT superfamilies, both *Copia* and *Ty-3* were 2.6 times more abundant in *P. edelbergii* than in *P. veris*. Despite differences in relative abundance of TE classes, *P. edelbergii* and *P. veris* were both enriched in TEs towards pericentromeric regions (**Figure 2a**; Potente et al., 2022a). When plotting the distribution of each TE superfamily separately, we determined that this pattern was largely driven by LTR-RTs, while other TE superfamilies were uniformly distributed across chromosomes (**Figures S6, S7**).

A likely explanation for the higher TE content of *P. edelbergii* vs *P. veris* is the smaller effective population size (N_e_) of the former, restricted to two small populations in Afghanistan, while the latter is distributed across Eurasia. Consequently, *P. edelbergii* should experience a relaxation in the efficacy purifying selection, resulting in less efficient purging, and consequent accumulation, of TEs (Le Rouzic & Deceliere, 2005). Additionally, *P. edelbergii* is self-compatible (Al Wadi & Richards, 1993) selfing could, at once, further decrease the efficacy of purifying selection and increase homozygosity, thus reducing the probability of TE removal (e.g. for LTR-RTs) via ectopic recombination (Montgomery et al., 1991).

### Gene annotation and expression

Using a combination of *ab initio*, evidence-based and comparative gene-prediction approaches, we identified 21,513 protein-coding genes in *P. edelbergii*. Three sources of evidence supported the high quality of our gene annotation. First, 21,022 gene models (97.71% of the total) were supported by RNA-seq evidence, i.e. were covered at least partially by RNA-seq reads, with 19,446 gene models (90.39% of the total) having RNA-seq coverage for their entire coding region (**Table S5**). Second, the ratio between mono-exonic and multi-exonic genes (mono:multi ratio) was 0.18, thus being very close to the value of 0.2 that has been proposed as optimal for angiosperm genomes (Jain et al., 2008; Vuruputoor et al., 2023). Third, the BUSCO completeness score of 95.5% obtained for the *P. edelbergii* proteome was the highest among all Ericales investigated here (**Figure S3, Table S2**).

The number of predicted genes in *P. edelbergii* (21,512) was substantially smaller than that of *P. veris*. (34,580). However, in *P. veris* only 64.48% of the genes were supported by RNA-seq evidence for their entire coding region, despite the availability of a larger RNA-seq data set for that species (Potente et al., 2022a). Additionally, the *P. veris* gene set was characterized by a slightly higher mono:multi ratio (0.21). Thus, the larger number of genes in *P. veris* compared to *P. edelbergii* is likely explained by a poorer gene annotation in *P. veris*, producing an excess of gene models.

The four *P. edelbergii* RNA-seq samples used for gene annotation were also used to quantify expression in different tissues. The total number of expressed genes was 19,272 (89.7% of the total), of which 16,251 (75.7%) were expressed in all samples (**Figure 3a**,**b**,**d**). Floral samples were characterized by a higher number of expressed genes compared to the leaf sample, and ‘flower’ and ‘young floral bud’ samples had the highest number of tissue-specific genes (**Figure 3b**,**c**). To investigate the tissue-specificity of expression, we calculated the coefficient of variation (CV = standard deviation/mean expression across samples) and the tau index for all genes across all samples. Both CV and tau distributions were skewed towards zero, further confirming that *P. edelbergii* genes are characterized by an overall low tissue-specific expression (**Figure 3e**,**f**).

### The *S*-locus translocated since the divergence between *P. edelbergii* and *P. veris*

The genome assemblies of two distantly related (18 Mya) heterostylous species that share the same origin of heterostyly (de Vos et al., 2014b) allowed us to ask for the first time whether the structure, composition, and sequence of the *S*-locus are maintained across macroevolutionary timescales. To address this question, we first searched for a region of the *P. edelbergii* genome syntenic to the region containing the *S*-locus in *P. veris*, then we checked whether such region of *P. edelbergii* contained genes homologous to the *P. veris S*-genes.

The region containing the *S*-locus in *P. veris*, located in chromosome 1 (position 27.43–27.70 Mb), is largely syntenic with a region on chromosome 6 of *P. edelbergii* (position 28.4-28.7 Mb) that however does not contain any *S*-gene, but only two tandemly-repeated copies of the *CFB* gene (**Figure 4a**). However, we identified orthologs for all *P. veris S*-genes, except *CCM*^*T*^, in a 633,644-kb (37,912,377-38,546,021 Mb) region of *P. edelbergii* chromosome 2 that was embedded in a region not collinear with *P. veris* but collinear with the outgroup *A. corniculatum* (**Figure 4b**). Our study thus shows that the *S*-locus of *P. edelbergii* extends over approximately 634 kb, which is 2.4 times the length of the *P. veris S*-locus (ca. 260 kb). Furthermore, *S*-genes are ordered and oriented differently in *P. edelbergii* (*CYP*^*T*^*-PUM*^*T*^*-GLO*^*T*^*-KFB*^*T*^) and *P. veris* (*GLO*^*T*^*-CYP*^*T*^*-PUM*^*T*^*-KFB*^*T*^; **Figure 4c**). These results suggest that the *S*-locus translocated and rearranged since the divergence between *P. veris* and *P. edelbergii*. However, the lack of highly-contiguous genome assemblies from more distantly related *Primula* species prevented us from disentangling which of the two species retained the ancestral *S*-locus genomic location and gene order. We observe that, even following translocation, the *S*-locus has retained its position near the centromere in the two species. This observation may indicate a possible evolutionary constraint, wherein the *S*-locus appears to necessitate to be located in a low-recombining region, such as the pericentromeric region, potentially to maintain linkage disequilibrium with one or more flanking genes.

We then asked whether the *S*-locus was hemizygous also in *P. edelbergii*, as found in all other heterostylous species studied to date (Fawcett et al., 2023; Gutiérrez-Valencia et al., 2022; Li et al., 2016; Shore et al., 2019; Yang et al., 2023; Zhao et al., 2023), by checking whether the *S*-locus region was characterized by halved sequencing depth, an approach commonly used to detect hemizygous supergenes (Gutiérrez-Valencia et al., 2021). The *S*-locus had approximately the same sequencing depth as the rest of chromosome 2 (mean *S*-locus depth: 154.6x; mean chromosome 2 depth: 156.1x; **Figure 4d**). A similar result was obtained when focusing only on sequencing depth of genic regions (mean depth for *S*-genes: 164.9x; mean depth for all genes on chromosome 2: 160.2x; **Figure S8**). These findings imply that the *S*-locus is present in both haplotypes of *P. edelbergii*. Finally, we identified no heterozygous SNPs in the *S*-locus, nor in the regions flanking it, as the closest heterozygous SNPs were located at positions 37,265,444 and 42,563,585, i.e. 647 kb upstream and 4.02 Mb downstream of the *S*-locus, respectively (**Figure 4d**). Therefore, the *S*-locus of the *P. edelbergii* thrum individual used for the present genome assembly is entirely homozygous, preventing the possibility that it might contain one dominant and one recessive allele.

The homozygosity of the *P. edelbergii S*-locus represents a deviation from all heterostylous *Primula* species studied so far, where the *S*-locus is hemizygous in thrums and absent in pins (Cocker et al., 2018; Li et al., 2016; Potente et al., 2022a). We propose two explanations to account for this result. A first possibility is that the genetic architecture of heterostyly in *P. edelbergii* differs from that observed in other heterostylous species, with the *S*-locus not being hemizygous in thrums. However, we find this explanation unlikely because pins and thrums occur in a 1:1 ratio in *P. edelbergii* wild populations (Richards, 2003), which would be impossible if all thrums had a homozygous *S*-locus. A second possible explanation is that the *P. edelbergii* individual used for the genome assembly represents a homozygous thrum occasionally arising from selfing of self-compatible hemizygous thrums, given that *P. edelbergii* is self-compatible (Al Wadi & Richards, 1993). We favor this possibility because this specific individual comes from the Botanical Garden München-Nymphenburg (Germany) and likely stems from one of the few plants introduced into cultivation in 1969 from the only two known populations of this species endemic to Afghanistan (Richards, 2003; see Materials and Methods). To disentangle between these two possibilities, one should sequence multiple pin and thrum individuals but, unfortunately, our impossibility of accessing wild populations precluded us from doing so.

The *S*-locus translocation reported here resembles the case of the strawberry *Fragaria virgininiana*, in which a supergene containing the sex-determining region (SDR) translocated three times in less than a million years as part of a ca. 25-kb hemizygous cassette (Tennessen et al., 2018). The scenario proposed to explain the multiple SDR translocations in *F. virginiana* was that the SDR expanded at each translocation, possibly adding genes under antagonistic selection that the various translocations kept in a hemizygous state (Tennessen et al., 2018). Such an explanation could fit the findings for *P. edelbergii* vs *P. veris* but the unavailability of *P. edelbergii* resequencing data precluded us from investigating whether any additional gene with putative antagonistic function is linked to the *S*-locus in this species.

### No major changes in *S*-genes can explain self-compatibility in *P. edelbergii*

Unlike most heterostylous *Primula* species (including *P. veris* and *P. vulgaris*), both floral morphs of *P. edelbergii* are self-compatible (Al Wadi & Richards, 1993). This is puzzling, because in the *Primula* species studied so far, the *CYP*^*T*^ *S*-gene controls both style length and recognition of self-pollen in thrums (Huu et al., 2016, 2022). The self-compatibility of *P. edelbergii* could thus be explained by mutations in one or more *S*-genes or changes in their expression. Alternatively, a change in a gene outside the *S*-locus could cause the lack of self-incompatibility in *P. edelbergii*.

To detect possible differences in the coding sequences of the *S*-genes, we aligned the coding sequences of the *P. edelbergii S*-genes against those of *P. veris* and *P. vulgaris*. All *S*-locus genes had the same overall structure (i.e. same number and approximate length of exons) in the three species, and did not have any early stop codons or frameshifts that could disrupt their functions. Nevertheless, we observed several non-synonymous substitutions in the *P. edelbergii* S-genes relative to their *P. veris* and *P. vulgaris* homologs (**Figure S9**). Of all *S*-genes, *PUM*^*T*^ was the one showing the highest number of substitutions between *P. edelbergii* and the other two species (203 mismatches and 44 indels over a 762-bp alignment; **Table S6**). Although this finding could imply the involvement of *PUM*^*T*^ in the genetic control of incompatibility, it more likely represents a signature of relaxed selection, as suggested also by its elevated π_N_/π_S_ (Mora-Carrera et al., 2023) and d_N_/d_S_ values (see below; **Figure 5g**,**h**).

**Figure 5:**
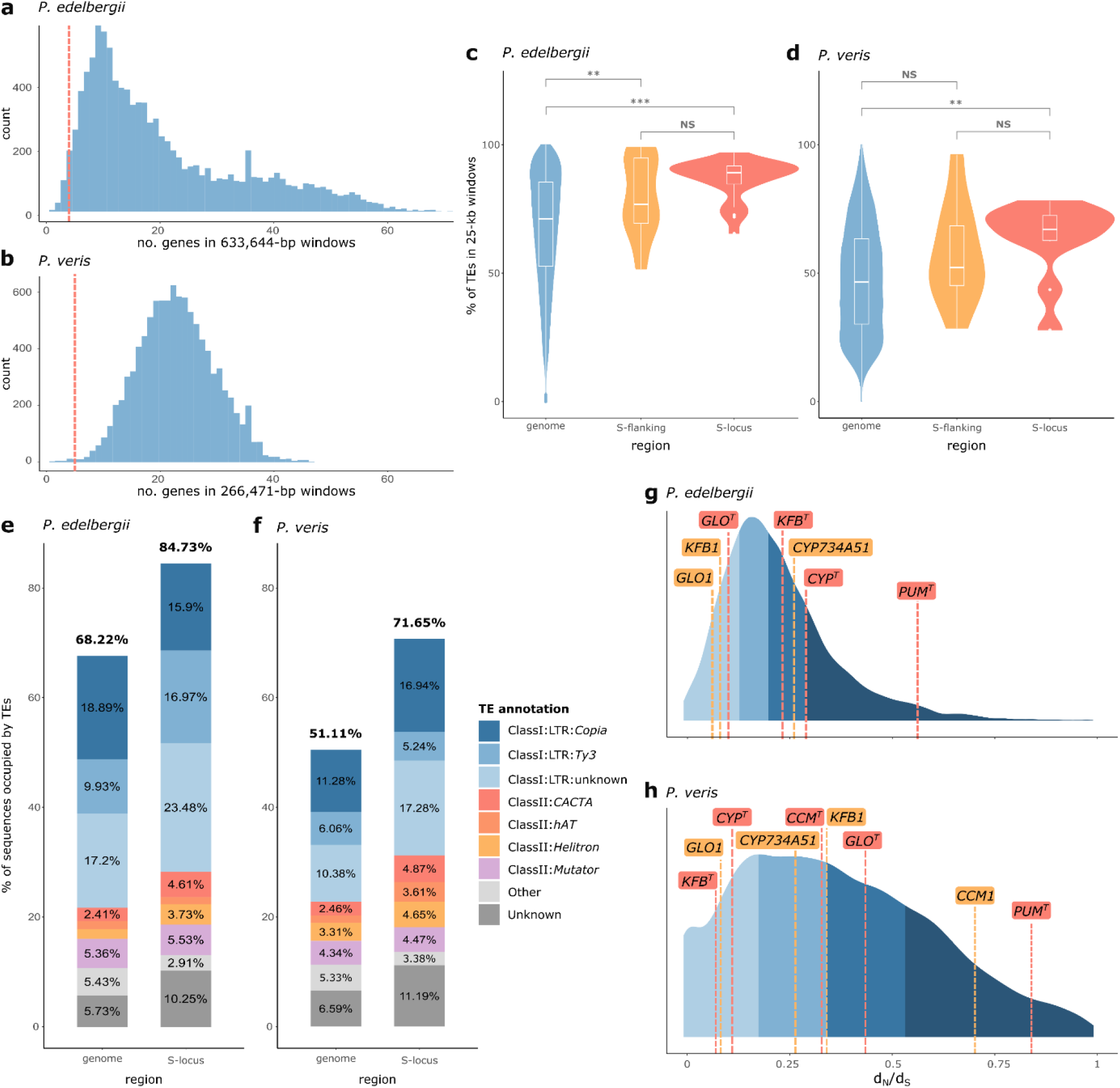
The *S*-locus does not show evidence of degeneration. (**a**,**b**) Distribution of the number of genes contained in genomic regions as long as the *S*-locus in *P. edelbergii* (a) and *P. veris* (b). The distribution was generated by randomly sampling 10,000 windows 633,644-bp long for *P. edelbergii* (a) and 266,471-bp long for *P. veris* (b). The vertical red dashed lines indicate the number of genes contained in the *S*-loci, i.e. four in *P. edelbergii* and five in *P. veris*. (**c, d**) Violin plots showing the proportion of sequence occupied by TEs, calculated in 25-kb non-overlapping windows for both *P. edelbergii* (c) and *P. veris* (d). Within each violin plot, the corresponding box plot is shown, with edges indicating the lower and upper quartiles, respectively, the middle line the median, and whiskers extending to 1.5 times beyond the upper and lower quartile. Red color refers to TE content in the *S*-locus, yellow to the regions immediately flanking the *S*-locus, and blue to the rest of the genome. ** 0.001 < p ≤ 0.01; *** p ≤ 0.001; NS non-significant; Wilcoxon rank-sum test. (**e, f**) Proportion of sequence occupied by TEs in the whole genome (left) and in the *S*-locus (right) of *P. edelbergii* (e) and *P. veris* (f), colored by superfamilies as indicated by the legend. To ease visualization, text indicating percentage for TEs occupying <2% of the sequence were removed; details are reported in Table S3. (**g, h**) Distribution of d_N_/d_S_ calculated between *P. edelbergii* and *P. vulgaris* (g) and between *P. veris* and *P. vulgaris* (h) for all genes in the genomes. Different shades of blue represent the four quartiles. The d_N_/d_S_ values for *S*-genes and their paralogs are indicated by red and yellow dashed lines, respectively.

To detect possible differences in the expression of *S*-genes, we compared the relative expression of each *S*-gene in the four samples for which we had generated RNA-seq data and found that all four *S*-genes were expressed in at least one developmental stage of the flower, i.e. the organ where they control heterostyly (**Figure S10**). Specifically, *CYP*^*T*^ and *GLO*^*T*^ had their highest expression in the ‘mature floral bud’ sample, while their expression was negligible in leaves; *PUM*^*T*^ was expressed at high level in both floral and leaf samples, confirming previous observations (Potente et al., 2022b; *KFB*^*T*^ expression was detected only in the ‘young floral bud’, corroborating the hypothesis that this gene is expressed only for a short temporal window (Potente et al., 2022b; **Figure S10**).

To summarize, our analyses did not detect any major change in the expression of any *S*-gene of *P. edelbergii* compared to self-incompatible *Primula* species. However, we found several differences in the coding sequences of virtually all *S*-genes, but could not determine which one, if any, was responsible for the decoupling of style elongation from self-incompatibility in *P. edelebergii*. It also remains a possibility that the lack of self-incompatibility in *P. edelbergii* might be caused by a change in a gene outside the *S*-locus, although such gene remains unknown. We note that the latter scenario could at once explain the occurrence of self-compatibility in both floral morphs, whereas a change in an *S*-gene would explain self-compatibility only in thrums.

### The *S*-locus is not degenerating in *Primula*

Two contrasting forces are expected to shape the evolution of hemizygous supergenes. On one hand, lack of recombination and reduced N_e_ (due to hemizygosity) are expected to decrease the efficiency of purifying selection, promoting genetic degeneration, i.e. increased gene loss and accumulation of deleterious mutations and TEs (Gutiérrez-Valencia et al., 2021). On the other hand, genes in a hemizygous supergene are expected to undergo stronger purifying selection than the genomic background because any deleterious mutation in them would be effectively dominant, as no homologous alleles are available to restore the wild-type phenotype (Gutiérrez-Valencia et al., 2021). Indeed, previous studies found no conclusive evidence of degeneration in the hemizygous *S*-loci of *P. veris* (Potente et al., 2022a), *Linum tenue* (Gutiérrez-Valencia et al., 2022), and *Nymphoides indica* (Yang et al., 2023).

We compared genetic degeneration in the *S*-loci of *P. veris*, known to be hemizygous (Li et al., 2016), and *P. edelbergii*, which we suspect to be hemizygous in wild populations (see above), and obtained similar results. In both species, we detected lower gene density in the *S*-locus than in the genomic background (**Figure 6a**,**b**; **Table S7**) and significantly higher TE content in the *S*-locus (*P. edelbergii*: 84.73%; *P. veris*: 71.5%) than the genome-wide average (*P. edelbergii*: 68.22%; *P. veris*: 51.11%) owing to an accumulation of both DNA and LTR transposons (**Figure 6e**,**f**). However, TE content within the *S*-locus did not exceed that of its genomic surroundings (**Figure 6c**,**d**), suggesting that either the elevated TE content of the *S*-locus does not stem from genetic degeneration caused by hemizygosity but rather from its pericentromeric location already characterized by high TE content, or recombination suppression extends over the borders of the *S*-locus. The lack of population genomic data prevented us from discerning between these two alternative hypotheses, but it is interesting to note that in *L. tenue*, in which the *S*-locus is not in a pericentromeric region, TEs were significantly higher in the *S*-locus than in its flanking regions (Gutiérrez-Valencia et al., 2022).

**Fig. 6:**
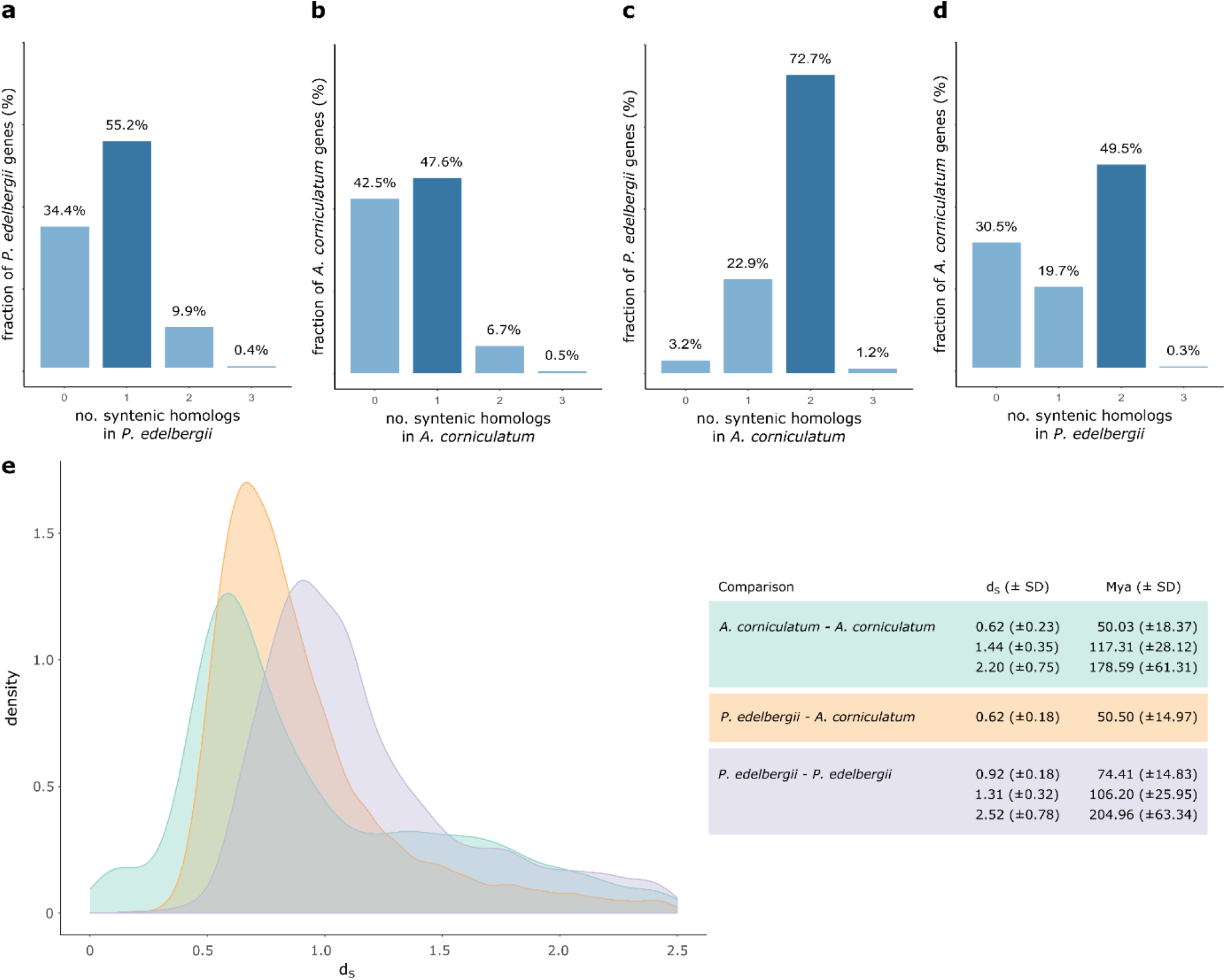
Are there one or two WGDs in Primulaceae? (**a, b**) Intragenomic syntenic depth for *P. edelbergii* (a) and *A. corniculatum* (b). Bar heights indicate the proportion of genes with zero to four paralogs in each genome. (**c, d**) Intergenomic syntenic depth between *P. edelbergii* and *A. corniculatum*. c) Proportion of *P. edelbergii* genes with different numbers of syntenic homologs in the *A. corniculatum* genome. d) Proportion of *A. corniculatum* genes with different numbers of syntenic homologs in the *P. edelbergii* genome. (**e**) Left: Distribution of d_S_ for paralogous gene pairs within *A. corniculatum* (turquois) and *P. edelbergii* (purple), in which each peak indicates a putative WGD, and for orthologous gene pairs between *A. corniculatum* and *P. edelbergii* (orange), whose peak indicates the divergence time between the two species. Right: table summarizing the statistically significant peaks identified in each d_S_ distribution by fitting a mixture of 1–3 normal distributions (for details, see Materials and Methods Figure S11), their respective standard deviation (SD) and age in million years (Mya).

We then searched whether the *S*-genes were accumulating more deleterious mutations than the genomic background by estimating the ratio of non-synonymous to synonymous substitutions (d_N_/d_S_) in *P. edelbergii* and *P. veris*. Again, similar results were observed in the two species. When compared to the genomic background, *PUM*^*T*^ was the only *S*-gene characterized by increased d_N_/d_S_, being 0.57 in *P. edelbergii* (97.19th percentile) and 0.85 in *P. veris* (88.48^th^ percentile). All other *S*-genes showed d_N_/d_S_ values within the first three quartiles of the d_N_/d_S_ distribution calculated for the entire genome (**Figure 5g**,**h**). This result indicates that *S*-genes do not accumulate deleterious mutations at a higher rate than the rest of the genome, with the possible exception of *PUM*^*T*^. Furthermore, the three *P. edelbergii S*-genes with a paralog (i.e., *GLO*^*T*^, *KFB*^*T*^, and *CYP*^*T*^) showed slightly higher d_N_/d_S_ values than their respective paralogs (*GLO1, KFB1*, and *CYP734A51*; **Figure 5g**), providing evidence for degenerative divergence from their closest gene copies. In *P. veris*, results were less clear, with *GLO*^*T*^ accumulating more deleterious mutations than its paralog *GLO1*, whereas all other *S*-genes had lower d_N_/d_S_ values than their respective paralogs (**Figure 5h**).

Taken together, our results imply that the lack of recombination in the *S*-loci of *P. edelbergii* and *P. veris* caused TEs accumulation without significantly affecting coding sequences, for *S*-genes did not accumulate more deleterious mutations than the rest of the genome. These results confirm previous conclusions in *Primula* and other genera that *S*-locus hemizygosity counteracts the detrimental effects in coding regions expected in non-recombining genomic regions, such as supergenes (Gutiérrez-Valencia et al., 2021, 2022; Potente et al., 2022a; Yang et al., 2023).

### Contrasting evidence on the polyploidization history of Primulaceae

Disentangling the evolutionary history of WGDs in Ericales has proved elusive (Larson et al., 2020; Nie et al., 2022). In Primulaceae, we previously identified a WGD (*Pv*-α) predating divergence of the closely related *P. veris* and *P. vulgaris* ca. 2.5 Mya (Potente et al., 2022a). However, the absence of high-quality genome assemblies for other Primulaceae prevented us from ascertaining whether *Pv*-α was specific to *Primula* and whether additional WGDs occurred in the family. The availability of the high-quality genome assembly of *P. edelbergii*, which diverged from the *P. veris-P. vulgaris* clade ca. 18 Mya (**Figure 1b**), together with the chromosome-scale assembly of another Primulaceae species, namely *A. corniculatum* (Ma et al., 2021), which diverged from *Primula* 46-53 Mya (The Angiosperm Phylogeny Group, 2016), allowed us to address these questions. To investigate the history of WGDs in Primulaceae we estimated intra- and inter-genomic syntenic depth for *P. edelbergii* and *A. corniculatum* and d_S_ within and between the two species.

In both *P. edelbergii* and *A. corniculatum*, the majority of genes (11,835 and 15,271, corresponding to 55.2% and 47.6% of the total gene content, respectively) had a single paralog in the genome (**Figure 6a**,**b**), suggesting that both *P. edelbergii* and *A. corniculatum* underwent a single, recent WGD. Such a conclusion was further supported by an interspecific syntenic depth analysis showing a 2:2 correlation between the two species, indicating that most *P. edelbergii* genes have two homologs in *A. corniculatum* and vice versa (**Figure 6c**,**d**).

To ascertain whether the polyploidization signatures identified in *P. edelbergii* and *A. corniculatum* correspond to a WGD shared between the two species or if each species underwent independent WGDs, we calculated d_S_ within and between *P. edelbergii* and *A. corniculatum* and identified statistically significant peaks (**Figure 6e, Figure S11**). The within-species d_S_ distribution of *P. edelbergii* showed three statistically significant peaks at 0.92, 1.31, and 2.52 (**Figure 6e**, purple distribution), corresponding to three previously described WGD events: *Pv-α* at 74.41 Mya; *Ad-β* at 106.2 Mya; *γ* triplication at 204.96 Mya, shared by all eudicots (Potente et al., 2022a). The within-species d_S_ distribution of *A. corniculatum* showed, in addition to two peaks corresponding to the *γ* triplication and the *Ad-β* WGD, at 2.20 (178.59 Mya) and 1.44 (117.31 Mya) respectively, a more recent peak at 0.61 (50.03 Mya) that appears to be too recent to correspond to *Pv-α*, hence possibly representing a newly discovered WGD (**Figure 6e**, turquois distribution). The d_S_ calculated between *P. edelbergii* and *A. corniculatum* peaked at 0.62 (50.50 Mya; **Figure 6e**, yellow distribution), suggesting that *Pv-α* preceded speciation while the putative WGD observed in *A. corniculatum* occurred after speciation. This result is puzzling because it contrasts the evidence we obtained with our previous analyses; if *Pv-α* preceded the divergence between *P. edelbergii* and *A. corniculatum*, and *A. corniculatum* underwent an additional WGD, we should observe: i) the majority of *A. corniculatum* genes having two paralogs instead of one (**Figure 6b**); ii) most *P. edelbergii* genes having three or four homologs in *A. corniculatum* instead of two (**Figure 6c**); iii) four significant peaks in the *A. corniculatum* d_S_ distribution, instead of three (**Figure 6e**, yellow distribution).

In summary, different analyses provided different results on the evolutionary history of WGDs in Primulaceae: while the intra- and inter-genomic syntenic depths between *P. edelbergii* and *A. corniculatum* suggest a single, shared WGD, the d_S_ analyses suggest that *A. corniculatum* underwent an additional WGD after the divergence from *Primula*. To disentangle the number and timing of WGDs in Primulaceae, more genome assemblies of closely related species will be required.

## Conclusions

We generated a chromosome-scale genome assembly of *P. edelbergii* using a combination of short- and long-read sequencing (**Figures 1**,**2a**). A whole-genome comparison between *P. edelbergii* and *P. veris* revealed several chromosomal rearrangements (**Figure 2b**) including the *S*-locus supergene, which ‘jumped’ to a new genomic location, being in chromosome 1 in *P. veris* and chromosome 2 in *P. edelbergii* (**Figure 4a**,**b**). Furthermore, the *S*-genes were reshuffled within the *S*-locus (**Figure 4c**) but this did not considerably change their expression pattern (**Figure 3, Figure S10**). We also found no signatures of degeneration in the *S*-loci of *P. edelbergii* and *P. veris*, confirming previous observations from other heterostylous taxa and showing that hemizygosity counteracts the genetic degeneration expected in non-recombining regions (**Figure 5**). Finally, we investigated the polyploidization history of Primulaceae, obtaining contrasting results on the number and timing of WGDs in this family (**Figure 6**).

Concluding, our study represents the first comparison of genomic architecture and sequence of *S*-loci from two distantly related species sharing the same origin of heterostyly providing, to our knowledge, the first evidence of translocation for a supergene not involved in sexual determination.

## Acknowledgements

This work was supported by the Swiss National Science Foundation to E.C. (Grant No. 175556) and the University Research Priority Program (URPP) *Evolution in Action* of the University of Zurich to N.Y. We thank Markus Meierhofer and Rayko Jonas for taking care of the plants in the glasshouse. We thank the Functional Genomics Center Zurich (FGCZ) staff for generating the Illumina sequences and their support throughout this project. We also thank the Botanischer Garten München-Nymphenburg (Munich, Germany) for providing *P. edelbergii* seeds through the International Plant Exchange Network (IPEN).

## Author Contributions

G.P., E.C., B.K., and N.Y. designed the study. B.K. and N.Y. collected seeds, cultivated *P. edelbergii* specimens, performed nucleic acid isolation and long-read DNA sequencing. G.P. and N.Y. performed the genome assembly. G.P. performed all downstream analyses, with help from N.Y., E.M.C., and P.S.. G.P. and E.C. wrote the manuscript, with input from all authors.

