## Supplementary Figures for "The *Primula edelbergii S*-locus is an example of a jumping supergene"

Figure S1: Genome-size estimate via k-mer analysis ... 2

Figure S2: Genome assembly quality evaluated by k-mer analysis ... 3

Figure S3: Genome and proteome evaluation by BUSCO ... 4

Figure S4: Dot-plots of putative centromeric regions ... 7

Figure S5: Synteny between *P. edelbergii* and *P. veris* ... 8

Figure S6: TE distribution across chromosomes in *P. edelbergii* ... 9

Figure S7: TE distribution across chromosomes in *P. veris* ... 10

Figure S8: Sequencing coverage on *S*-genes and flanking genes ... 11

Figure S9: Protein alignment of *S*-genes ... 13

Figure S10: Expression profile of *S*-genes and their paralogs in *P. edelbergii* ... 14

Figure S11: $d_S$ distributions used for dating WGDs ... 15

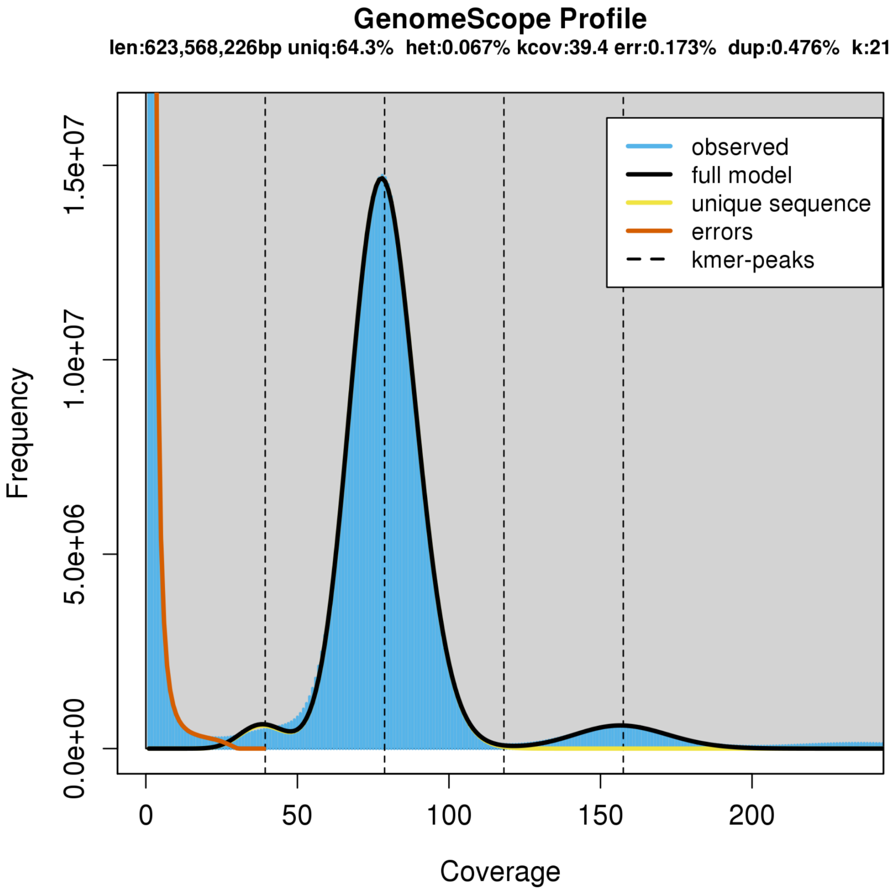


**Figure S1: Genome-size estimate via k-mer analysis**

The *P. edelbergii* genome size estimated using 21-mers corresponded to 623,568,226 bp. The slight underestimate, compared to the genome-assembly size (671.82 Mb), might be due to an underestimate of repetitive regions by GenomeScope. Indeed, GenomeScope estimated repetitive elements to comprise 400.90 Mb, while RepeatMasker identified 452.56 Mb of repetitive elements in the assembly.

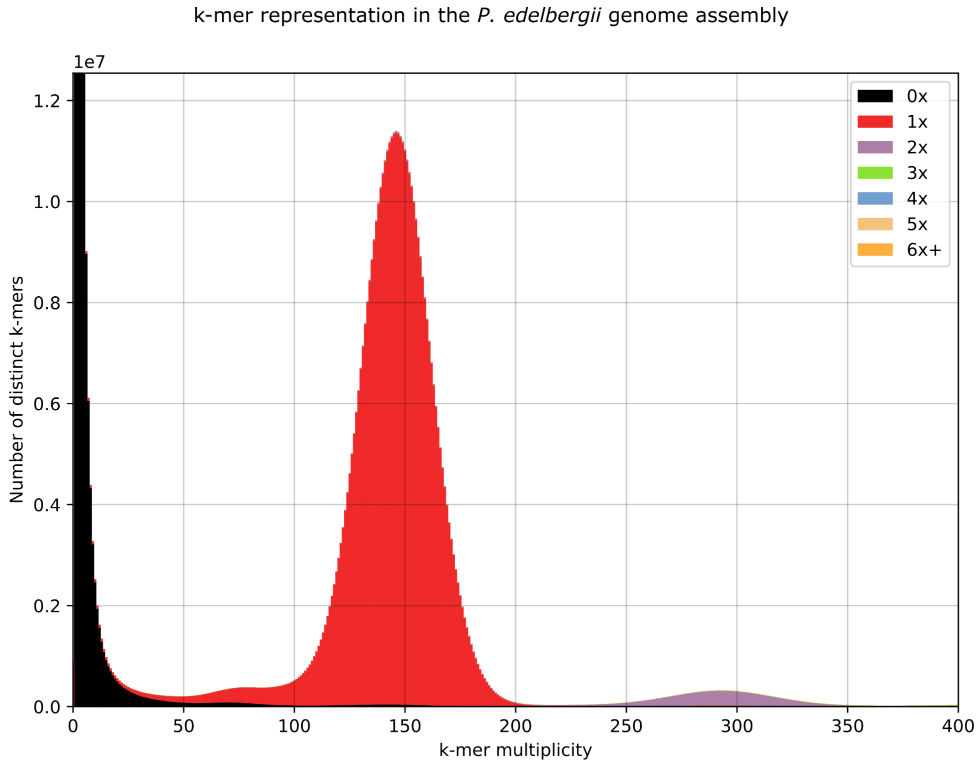


**Figure S2: Genome assembly quality evaluated by k-mer analysis**
The histogram represents the k-mer spectrum (k-mer frequency vs number of distinct k-mers) obtained with the K-mer Analysis Toolkit (KAT) using illumina reads. Compared to the GenomeScope k-mer plot (Figure S1), here each bar of the histogram is colored based on how many times each k-mer is present in the genome assembly. As expected, k-mers with very low multiplicity, thus representing sequencing errors, are not present in the assembly (black). The high homozygosity of the genome is reflected by the red peak at k-mer multiplicity ~150 being much higher than the heterozygous peak (at k-mer multiplicity ~75). A third short peak (at k-mer multiplicity ~300) represents genomic regions that are duplicated in the genome and as expected contains k-mers present twice in the assembly (purple).

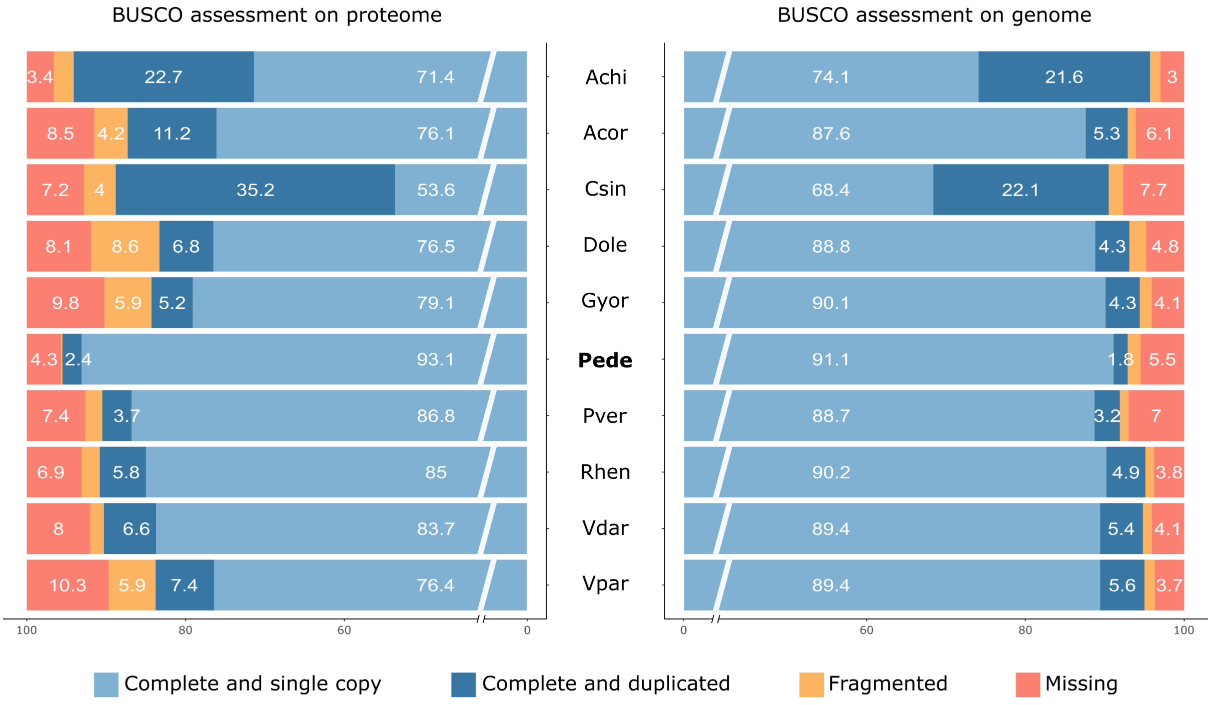


**Figure S3: Genome and proteome evaluation by BUSCO**

Completeness BUSCO scores for ten Ericales proteomes (left) and genome assemblies (right), obtained using the eudicots_odb10 database, which contains 2,326 single-copy orthologs. Numbers in each bar represent the percentage of genes in each category. To ease visualization, percentages are not reported for fragmented genes that accounted for less than 3% of the total. The total number of complete BUSCO genes for each species is given by the sum of "Complete and single copy" (light blue) and "Complete and duplicated" (dark blue). The species included (and their respective abbreviations) are: *Actinidia chinensis* (Achi), *Aegiceras corniculatum* (Acor), *Camellia sinensis* (Csin), *Diospyros oleifera* (Dole), *Gilia yorkii* (Gyor), *Primula edelbergii* (Pede), *Primula veris* (Pver), *Rhododendron henanense* (Rhen), *Vaccinium darrowii* (Vdar), *Vitellaria paradoxa* (Vpar).

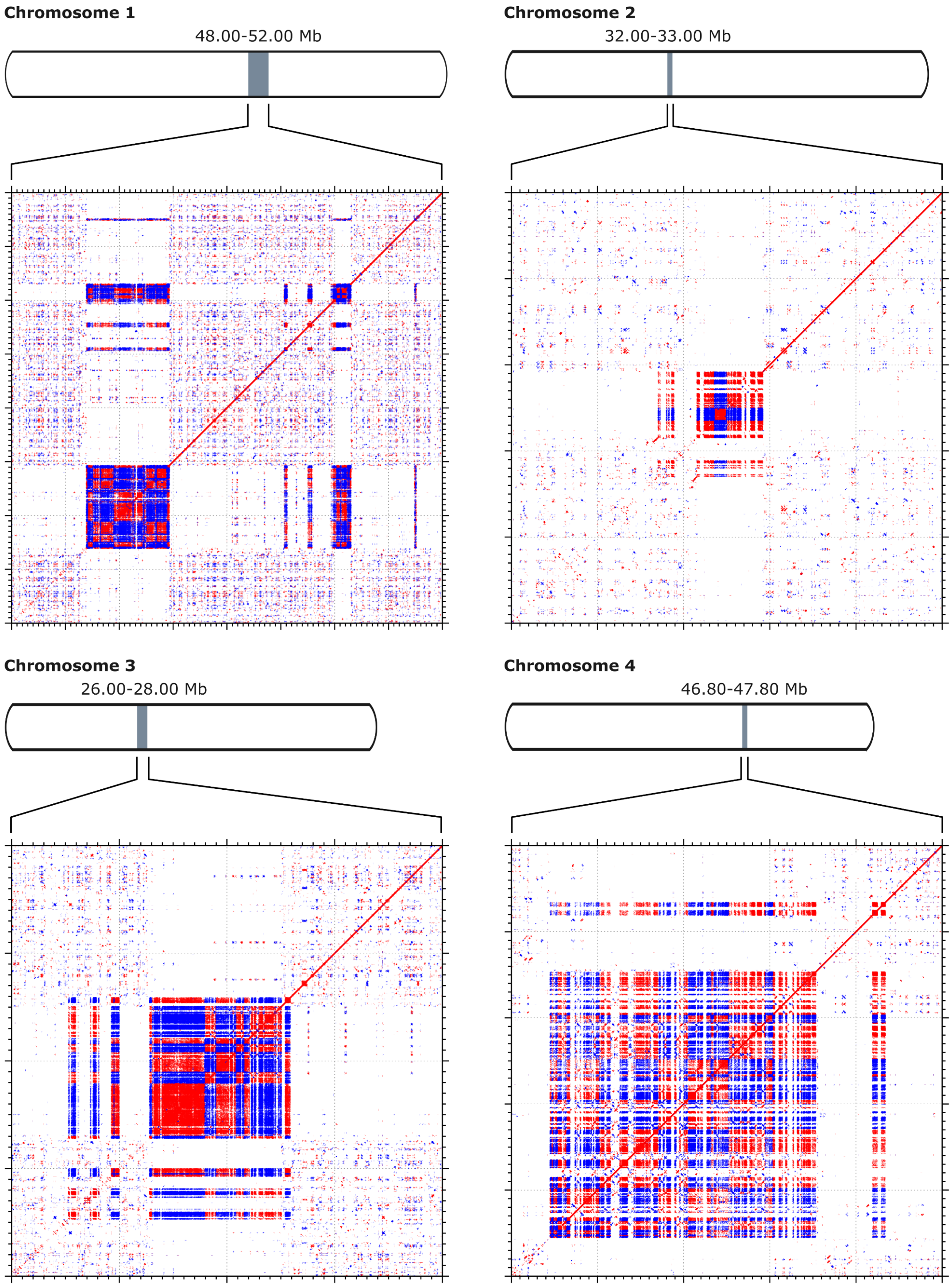

Chromosome 1
48.00-52.00 Mb
Chromosome 2
32.00-33.00 Mb
Chromosome 3
26.00-28.00 Mb
Chromosome 4
46.80-47.80 Mb

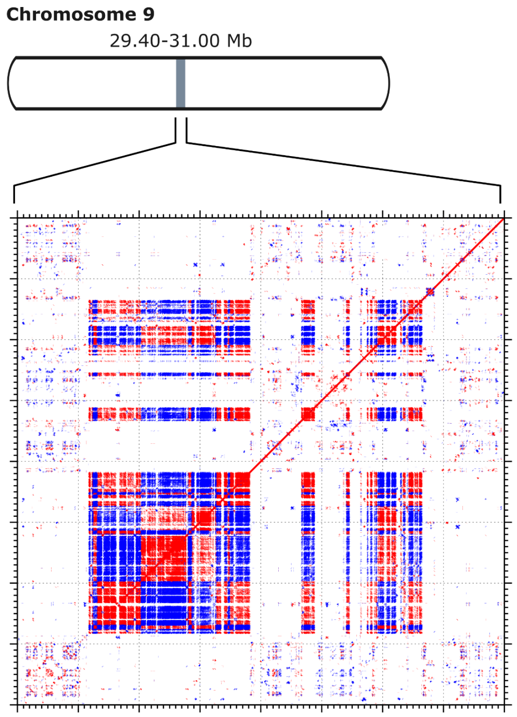


**Figure S4: Dot-plots of putative centromeric regions**
For each of the nine chromosomes of the *P. edelbergii* genome the following are shown: a schematic visualization of the chromosome with a grey bar indicating where the putative centromere is located (top), and a dot-plot of the putative centromeric region against itself (bottom), with red segments indicating regions collinear on the same strand and blue segments indicating regions collinear on the reverse complement. Dot-plots were generated with the MAFFT online service (mafft.cbrc.jp/alignment/server/index.html) using default parameters.

**a**

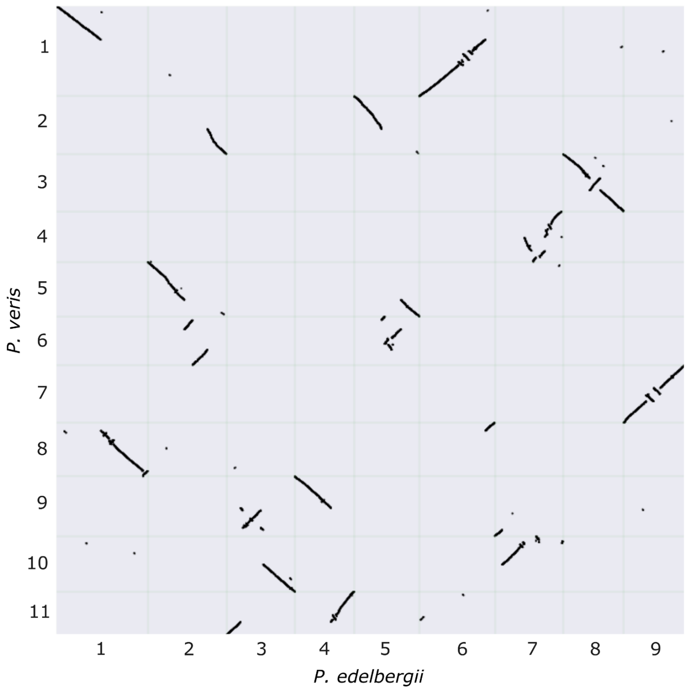


**b**

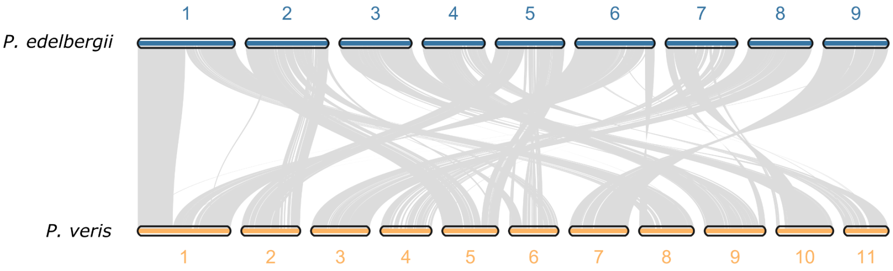


**Figure S5: Synteny between *P. edelbergii* and *P. veris***

**a.** Dotplot obtained by aligning the *P. edelbergii* genome assembly against the *P. veris* assembly; syntenic regions containing >5 collinear genes are represented by black marks. **b.** Synteny plot between *P. edelbergii* and *P. veris*; regions containing >5 collinear genes are connected by gray ribbons.

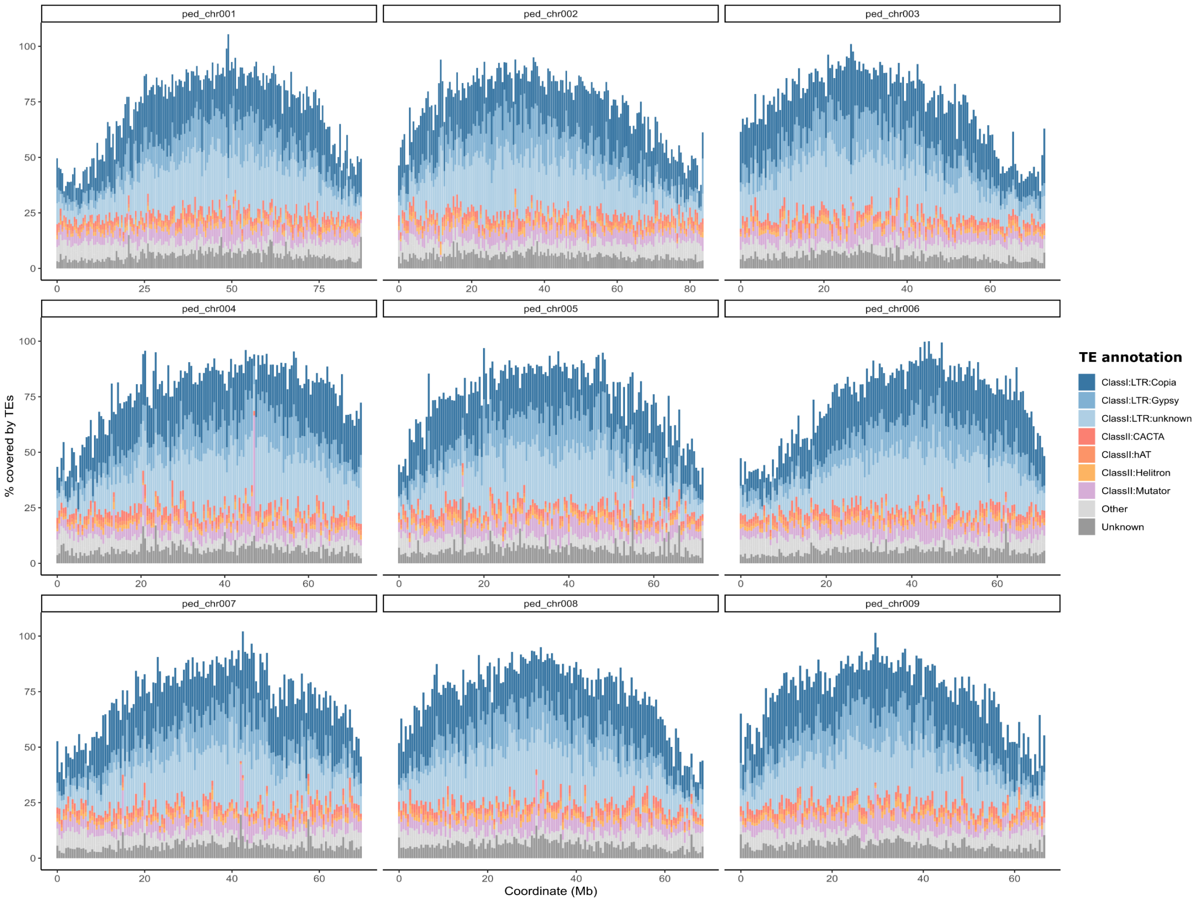


**Figure S6: TE distribution across chromosomes in *P. edelbergii***

Distribution of different TEs calculated in 500-kb windows across the nine chromosomes of *P. edelbergii*. Different TE families are colored as in the legend.

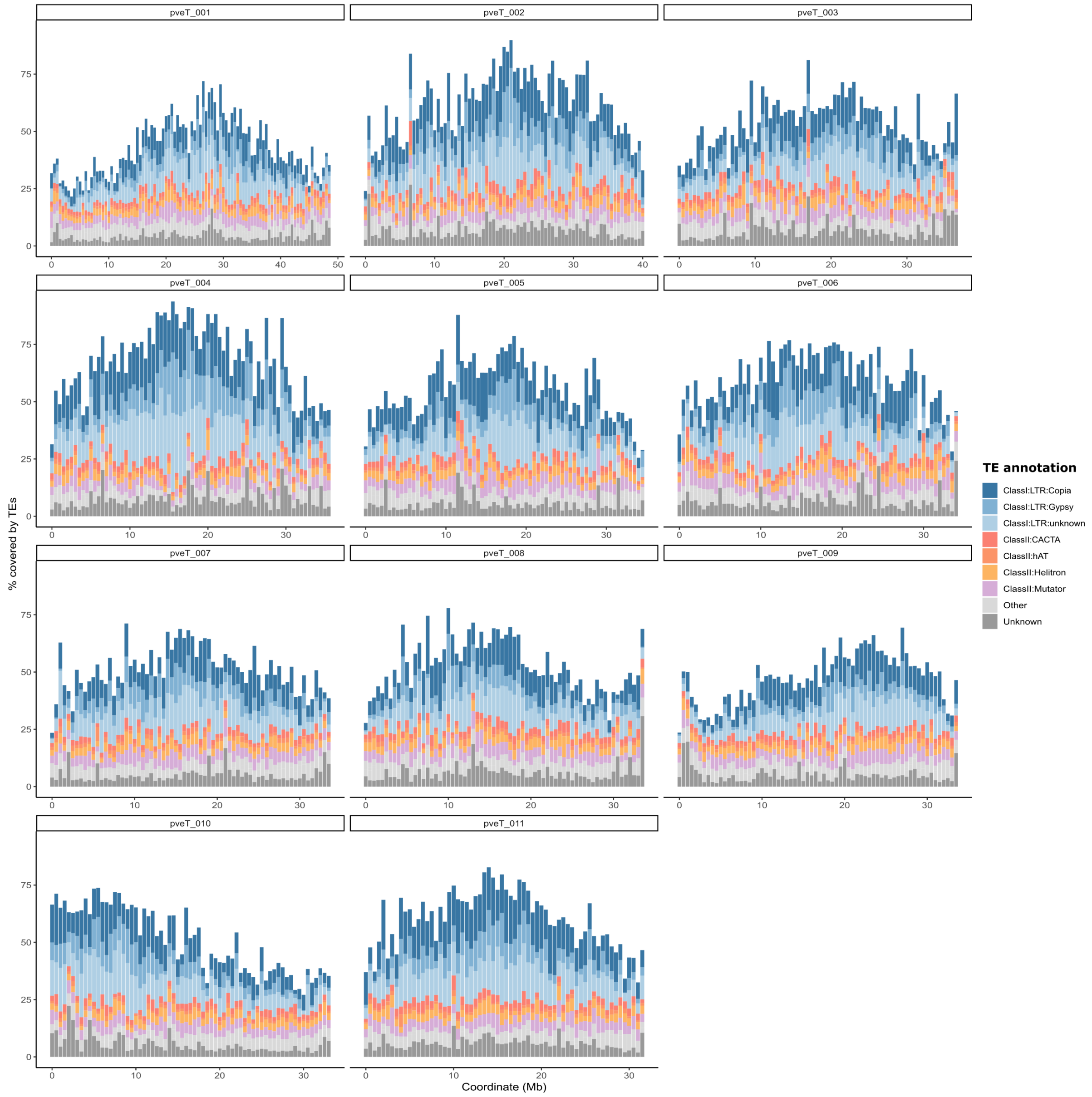


**Figure S7: TE distribution across chromosomes in *P. veris***

Distribution of different TEs calculated in 500-kb windows across the 11 chromosomes of *P. veris*. Different TE families are colored as in the legend.

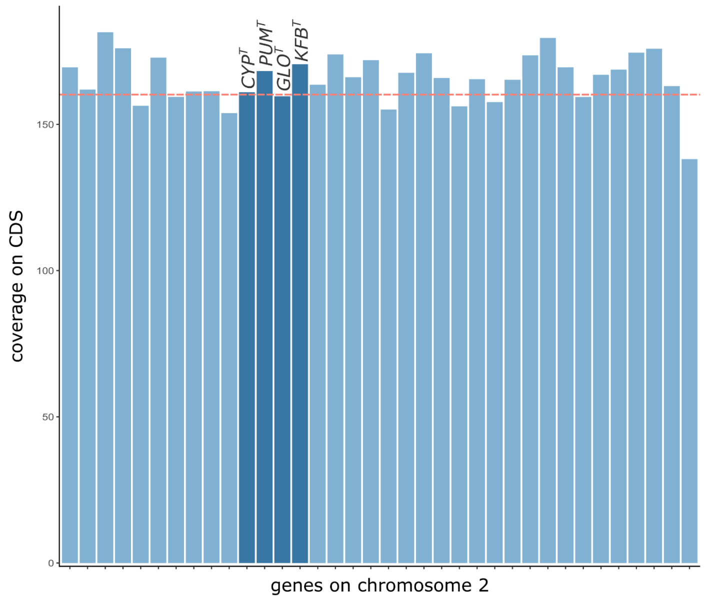


**Figure S8: Sequencing coverage on *S*-genes and flanking genes**
Sequencing coverage calculated on coding sequences for the four *S*-locus genes (dark blue) and the other 32 genes (light blue) contained between positions 37 Mb and 39.5 Mb on chromosome 2 (the same coordinates as in Figure 5d). A red dashed line indicates the mean per-gene coverage for all genes in chromosome 2 (n = 2,646; mean coverage = 160.2x), which does not differ from the one calculated for *S*-genes.

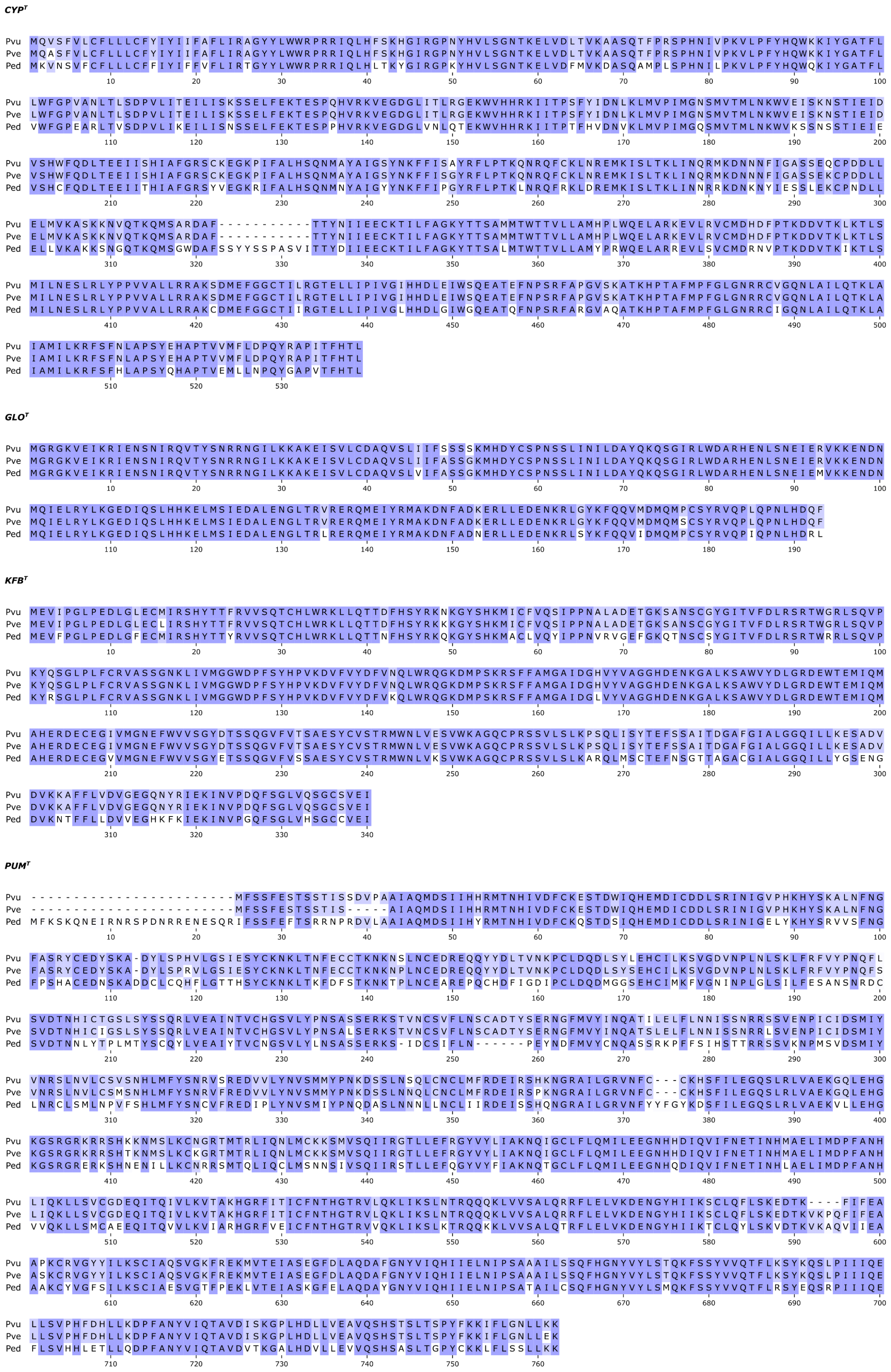
CYP^T
GLO^T
KFB^T
PUM^T
Pvu
Pve
Ped

**Figure S9: Protein alignment of *S*-genes**

Alignment of the aminoacidic sequences of the four *S*-genes as found in *P. vulgaris* (Pvu), *P. veris* (Pve), and *P. edelbergii* (Ped). For each sequence, sites with two matches are highlighted in dark purple, sites with one match are highlighted in light purple, and sites with no matches are left white.

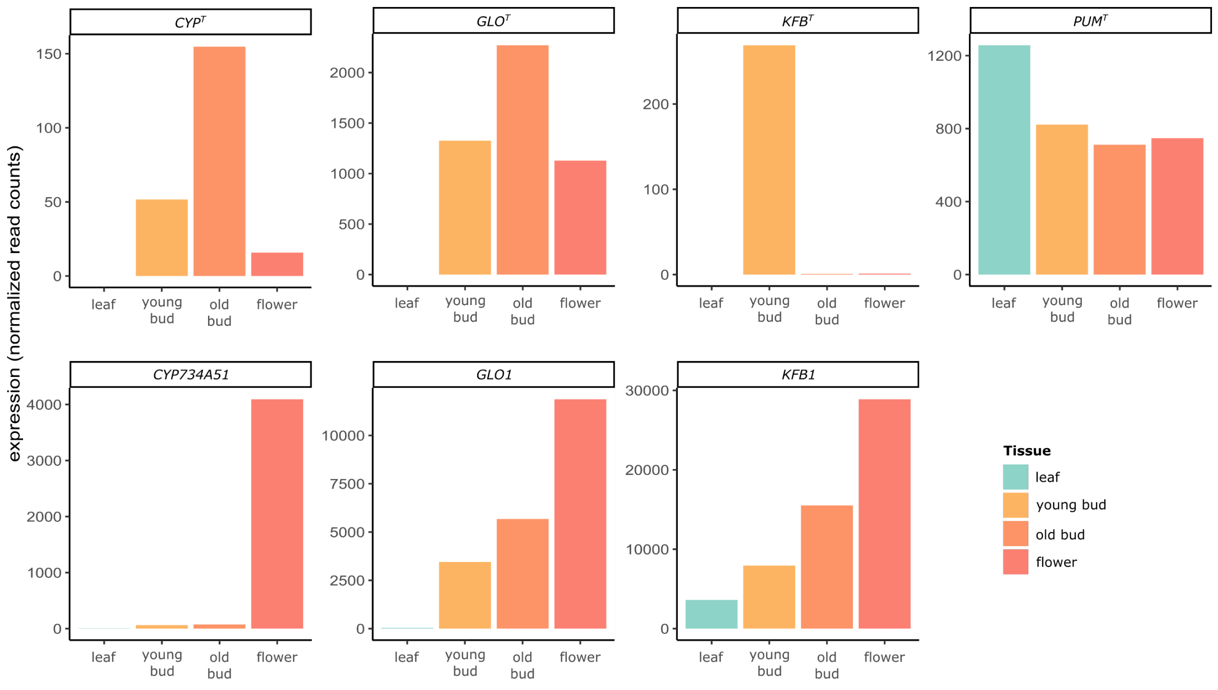


**Figure S10: Expression profile of *S*-genes and their paralogs in *P. edelbergii***
Barplots showing the number of normalized RNA-seq reads per each *S*-gene (top) and their respective paralogs (bottom) in the four tissues analyzed here. Each tissue is colored as in the legend.

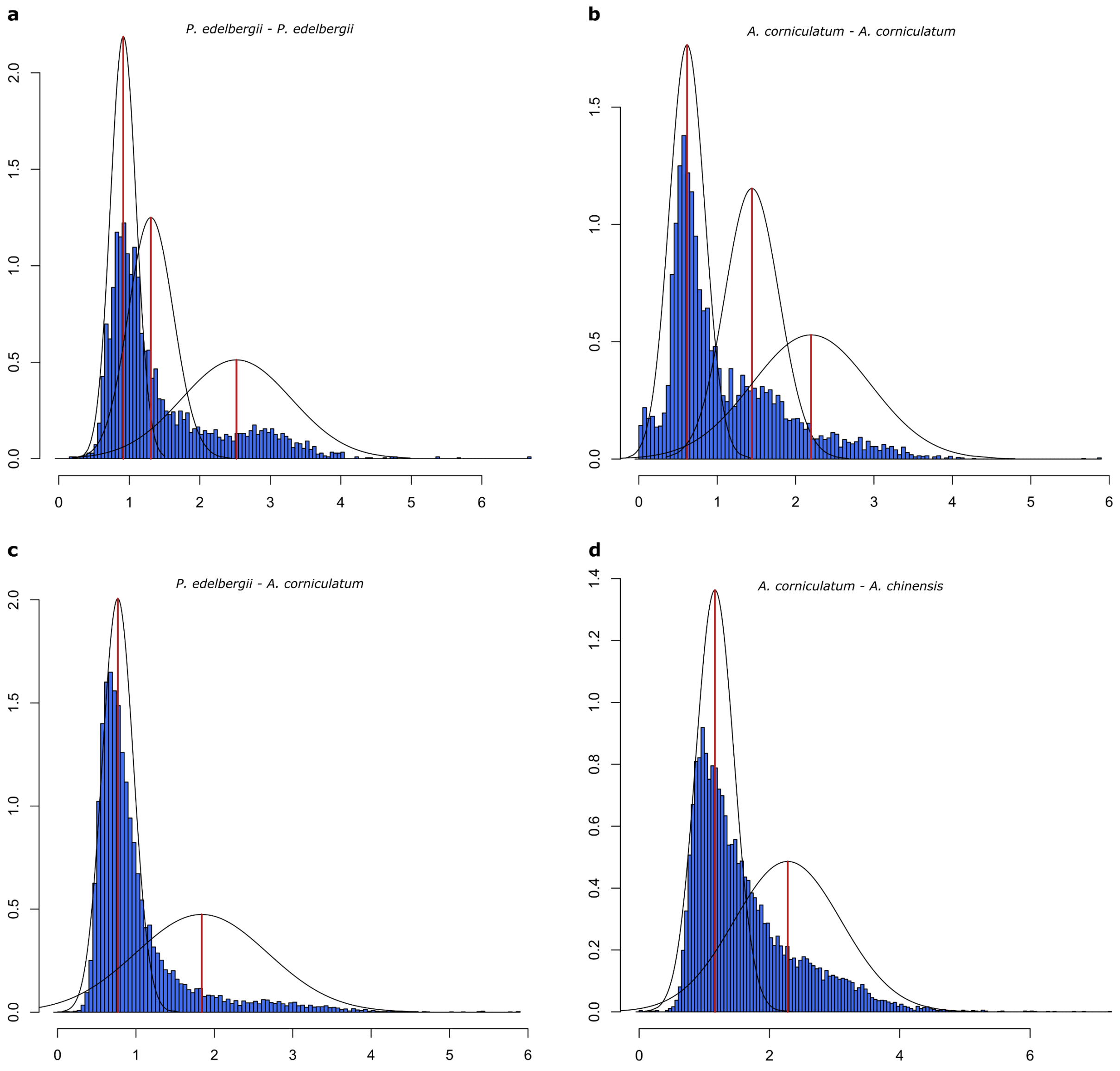


**Figure S11: $d_S$ distributions used for dating WGDs**

Histograms representing the $d_S$ distributions calculated for paralogous gene pairs in *P. edelbergii* (**a**) and *A. corniculatum* (**b**), and for orthologous gene pairs between *P. edelbergii* and *A. corniculatum* (**c**) and between *A. corniculatum* and *A. chinensis* (**d**). Black lines represent normal distributions corresponding to statistically significant peaks, with modal values shown as vertical red lines.
